# The transcriptome dynamics of single cells during the cell cycle

**DOI:** 10.1101/2019.12.23.887570

**Authors:** Daniel Schwabe, Sara Formichetti, Jan Philipp Junker, Martin Falcke, Nikolaus Rajewsky

## Abstract

Despite advances in single-cell data analysis, the dynamics and topology of the cell cycle in high-dimensional gene expression space remains largely unknown. Here, we use a linear analysis of transcriptome data to reveal that cells move along a circular trajectory in transcriptome space during the cell cycle. This movement occurs largely independently from other cellular processes. Non-cycling gene expression (changes in environment or epigenetic state) adds a third dimension and causes helical motion on a hollow cylinder. The circular trajectory shape indicates minimal acceleration of transcription, i.e. the cell cycle has evolved to minimize changes of transcriptional activity and its entailing regulatory effort. Thus, we uncover a general design principle of the cell cycle that may be of relevance to many other cellular differentiation processes.

**One Sentence Summary:** Cells traverse high-dimensional gene expression space in a 2D circular motion, thus minimizing changes of expression changes (“Acceleration”).

## Main Text

The cell cycle is a shared general principle of life, and core aspects of the cell cycle are conserved across eukaryotes (*1, 2*). However, as cell division rates vary massively across species and cell types, the cell cycle also needs to be plastic and coupled to cellular physiology. Despite a multitude of mechanistic studies, the topology of the cell cycle in gene expression space, as well as its degree of coupling to other cellular processes, remains largely unclear (*3–5*). For instance, it is not known if the cell cycle can be described as a two-dimensional oscillator or whether additional dimensions are needed, and it is not known which (if any) optimality principles govern gene expression changes along the cell cycle. In recent years, pseudo-temporal ordering of single cell transcriptomes has emerged as a powerful method for reconstruction of low-dimensional cell differentiation trajectories from high-dimensional single-cell RNA-seq data (*6*). The progression of a cell through the cell cycle can be represented as a trajectory in transcriptome space. Here, we examine transcriptomic snapshots of populations of asynchronous cycling cells in order to reconstruct, quantify and interpret the cell cycle as a dynamical system so as to define this trajectory as well as its underlying design principles.

We expect the trajectory to describe a periodic motion, completed once each time a cell divides. We further anticipate the trajectory to be constrained to a subspace with much lower dimension than the transcriptome space (~20,000 dimensions) because only a subset of genes is involved in the cell cycle and genes are known to interact in a highly coordinated manner (*1, 2*), i.e. groups of genes controlled by transcription factors and chromatin state are up- or downregulated together during the cycle (*7*). A priori, the transcriptomic trajectory describing the cell cycle might be a simple circle embedded in a plane, it might be wound up on a donut-like structure (torus), twisted and looped like a roller coaster in three dimensions or be even more complex in higher dimensions (Fig. 1A, Methods). The regulatory effort required to complete the cell cycle is closely related to the shape of the trajectory in transcriptome space (as explained in Methods).

**Fig. 1.**
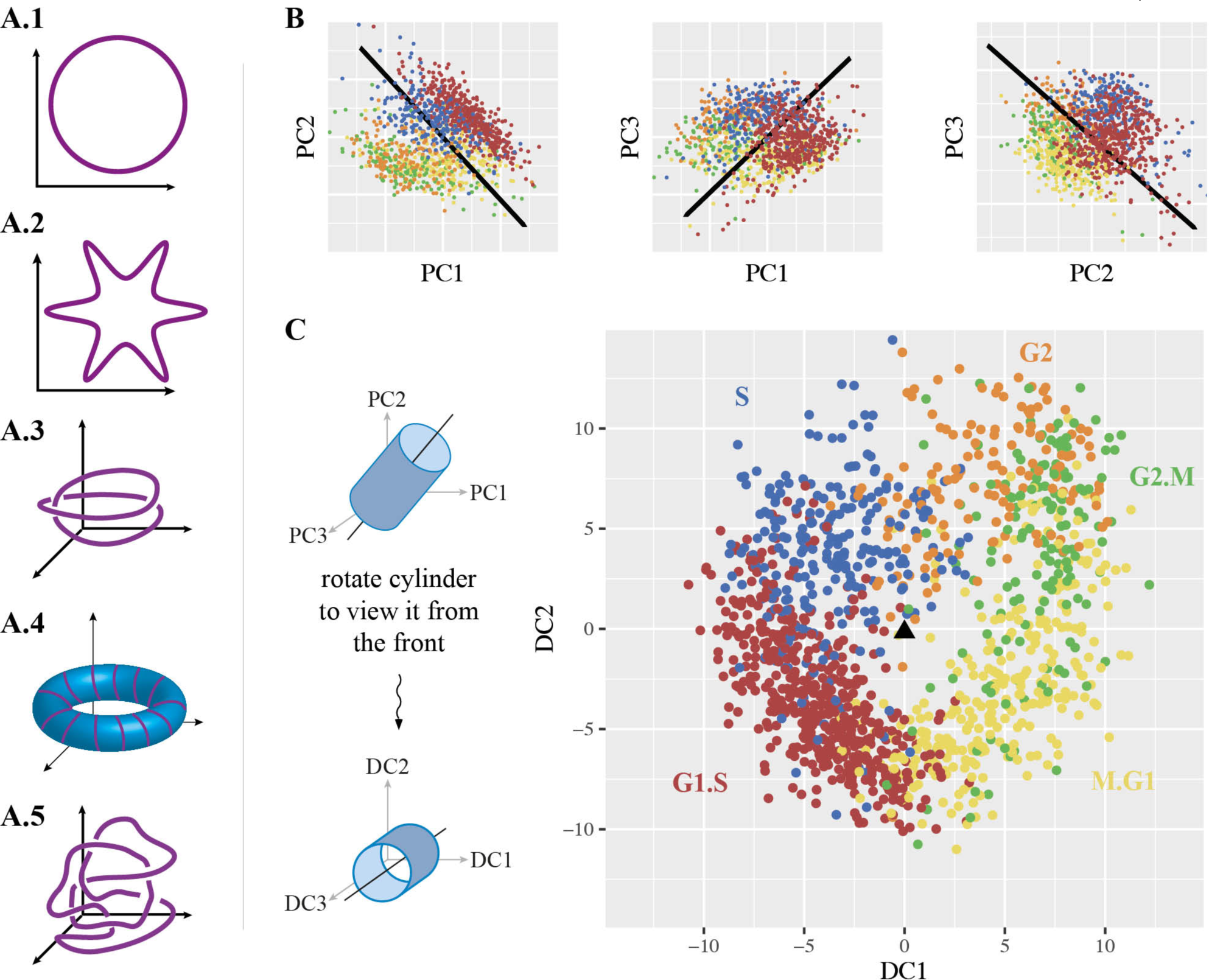
The cell cycle forms an annulus in two dimensions and a cylinder in three dimensions of transcriptome space. (**A**) Toy examples of possible shapes of cell cycle trajectories in transcriptome space. (**A.1**) A circle in two dimensions. (**A.2**) A star. (**A.3**) A cyclic trajectory requiring three dimensions with an upper and a lower loop. (**A.4**) A torus. (**A.5**) A three-dimensional motion comparable to a roller coaster. (**B**) Two-dimensional representations of the PC scores of all individual cells w.r.t. PC1, PC2 and PC3 exhibit clustering with regard to their computationally inferred cell cycle phases. A clear cyclic structure is absent. In three dimensions, the data form a slanted cylinder (cylinder axis indicated by black line). (**C**) If we rotate the cylinder and view it from the top or base, a clear cell cycle structure becomes visible. The rotated PCs are simple linear combinations of the original PCs, and are the dynamical components DC1 and DC2. The black triangle in the origin indicates the cylinder axis viewed from the front.

Due to cell-to-cell variability, cell cycle trajectories of individual cells, even of the same cell type, will not be identical and aligned. The collection of trajectories from a population of cells can be imagined as a tube in transcriptome space encompassing all trajectories. This tube may be called a manifold, and the volume of this manifold contains information on cell variability. We first set out to formally define the cell cycle manifold and then to identify trajectories within it with an RNA velocity analysis.

A HeLaS3 cell line was grown asynchronously and single-cell RNA sequenced deeply using an in-house optimized version of the Drop-seq protocol (*8, 9*) (Methods). The data set contains single-cell data of 1348 cells with a mean depth of almost 15,000 UMIs per cell (fig. S1A). We computationally inferred a cell cycle phase for each single cell by correlating its transcriptome data to known marker genes for different parts of the cell cycle (in particular G1.S, S, G2, G2.M and M.G1) (*8, 10*) (Methods). We restricted the data to 1536 detected highly variable genes. Furthermore, we transformed the data by calculating concentrations, multiplying by a scaling factor, log-transforming it and normalizing it across all genes and all cells (Methods) (*11*). While there is a large number of tools for pseudo temporal ordering of single cell transcriptomes (e.g. reCAT (*5*), Monocle (*12*), Wanderlust (*13*), Wishbone (*14*), PAGA (*15*)), these mostly specialize in non-linear manifold learning approaches. Here, however, we show that a fully linear approach is sufficient to isolate the cell cycle, which substantially facilitates downstream analysis and interpretation by preserving the geometrical structure of the trajectory.

After applying principal component analysis (PCA) to our data, we observed that the first three principal components (PCs) exhibit cell clustering according to the computationally inferred cell cycle phases (Fig. 1B). Additional PCs do not reflect the cell cycle (fig. S1B). None of the two-dimensional projections in PC space exhibits a clear periodic trajectory (Fig. 1B), suggesting that at least three PC-dimensions are necessary to define the cell cycle manifold.

Considering the space spanned by the first three principal components, we observed that the cloud of cells has the shape of a hollow cylinder which is slanted with respect to PC axes. By rotating the cylinder, we realized that viewing it from the top or base yields an annular shape in two dimensions that provides a surprisingly good representation of the cell cycle (Fig. 1C). Most notably, we obtained an exceptionally clear clustering and progression of cell cycle phases, which displays the expected order G1-S-G2-M. Additionally, we observed an area around the origin almost devoid of data points, in agreement with the principle that cells cannot skip phases. When viewed from the appropriate angle, the cell cycle is in fact contained in a two-dimensional plane.

The change of angle of view is achieved by a basic linear rotation of the PC space (Methods). The newly found axes after rotation – which we refer to as dynamical components (DC1 and DC2) – are linear combinations of the PCs involved in the rotation (Methods). The general steps of our algorithm REVELIO (REVEaling the cell cycle with a LInear Operator) from raw data to the two-dimensional cell cycle are outlined in fig. S2.

We were able to reproduce the results with other cell lines, including an additional HeLaS3, a HEK293 and a 3T3 data set where the form of the cell cycle varies from an annulus to a disc (fig. S3,S4,S5). We also managed to isolate the cell cycle into an annulus in just two dimensions even when utilizing all genes detected during sequencing (~10,000 genes) instead of only the highly variable genes (fig. S6). Hence, inclusion of additional genes into the analysis, which typically increases noise levels, does not alter the characteristics of the outcome. The limiting factor appears to rather be the sequencing depth because the less deeply sequenced a data set is, the more noise is incorporated due to e.g. increased amounts of dropouts. We hypothesize that in that case essential cell cycle information is missing for some cells, causing a positioning of cells near the origin. However, cells nearby the origin of the trajectory are biologically implausible as these cells would reside simultaneously in all cell cycle phases (we ruled out that these cells are simply dead cells by using standard apoptosis markers).

Overall, these results suggest that each individual cell describes a circular motion in transcriptome space, and due to cell-to-cell variability the collection of all trajectories describes an annulus-shaped manifold in two dimensions, or a hollow cylinder when considering three dimensions. In summary, we have revealed two remarkable design principles. (a) Low dimensionality: only two dimensions of the ~20,000 dimensional gene expression space are used for the cell cycle (in fact the lowest possible number of dimensions), and (b) circularity (the simplest and smoothest possible trajectory).

To further verify that the two-dimensional annulus does in fact represent the cell cycle, we investigated a number of characteristics starting with GO terms relating genes to function (*16, 17*). We found that a GO term analysis of the first three original PCs showed clear dominance of the cell cycle. However, only the two dynamical components DC1 and DC2 that create the annulus are heavily involved in the cell cycle, while the third dimension (DC3, parallel to the cylinder axis) does not contain any cell cycle related GO terms (Table S1). This supports the conclusion that we functionally isolated the cell cycle into two dimensions by a simple rotation. In agreement with this result, we found that DC3 is almost devoid of clustering with respect to the cell cycle phases (fig. S1C). These observations also apply to all additional dimensions of PC space.

Due to the simplicity of the obtained shape of the cell cycle, ordering the cells by their angle in a clockwise motion around the origin of the DC1-DC2-plane corresponds to the temporal order of the cell’s progression through the cell cycle. We divided the cycle into bins with equal cell numbers in order to investigate the development of total UMI counts per cell along the cycle. A sharp drop of average total UMI counts per cell by approximately a factor 1/2 occurs between the last and first bin of the cycle, at the overlap of G2.M and M.G1 cells (Fig. 2A). This is where cell division happens, and the drop of average total UMI counts strongly supports the temporal order of the cells derived by our algorithm. Measurements with other cell populations and other choices of bin sizes exhibit a similar drop of total UMI counts (fig. S3E, S4E, S5E, S6E, S7).

**Fig. 2.**
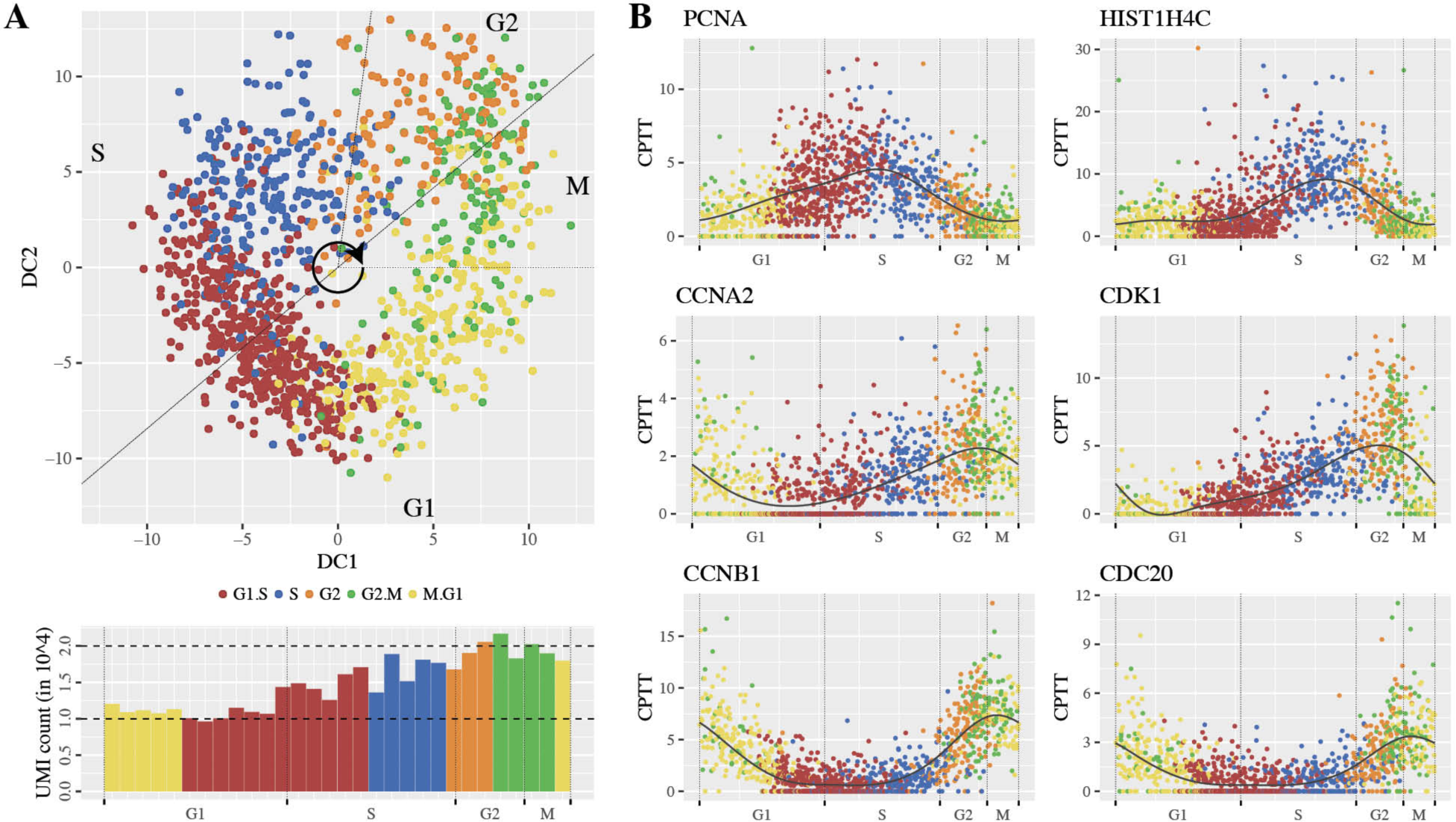
Drop of total UMI count by factor 1/2 at time of cell division and progression of individual genes confirm that the two-dimensional annulus corresponds to the cell cycle. (**A**) We dissect the cell cycle (top) into 30 bins containing equal amounts of cells and calculate the average total UMI count per cell for each bin. The sequence (bottom) yields a clear drop by almost factor 1/2 at the end of the green (G2.M) and the beginning of the yellow (M.G1) cluster. This is where mitosis is suspected to take place. (**B**) Extracting the angular motion of cells around the cell cycle, provides an order in time. We depict six time courses for well-known oscillating genes. For each individual gene, we relate the temporal order on the x-axis (cells are distributed evenly along the axis) to the distribution of the specific gene across all cells in counts per ten thousand (CPTT) (see Methods). This yields the dynamic behavior during the cycle from snapshot data. A spline of mean values is shown in black for each gene.

We observed that well-known cell cycle regulators peak at the expected time point along the cycle (Fig. 2B). For example, PCNA expression (activated during the G1-S transition (*2*)) rises during the end of G1 and beginning of S phase. The HIST1H4C gene exhibits a peak towards the end of S phase. Moreover, the peak transcription of CCNA2 during G2 phase falls in line with its role in guiding the cell through the S-G2 transition (*2*). The rising levels of CDK1 and CCNB1 indicate the formation of the mitosis-promoting factor, which pushes the cell into and through mitosis (*2*), hence they are expected to reach their maximum at the G2-M transition, just as we observe. Increased transcription of CDC20 during M phase occurs due to a negative feedback loop involving CDC20 which leads to the degradation of cyclin B causing the cell to exit mitosis (*2*). Time courses for other highly variable genes in our dataset strongly overlap with *Cyclebase* (*18*) (data not shown), confirming that the annulus and the implied temporal order of cells correspond to the cell cycle.

The fact that we find the cell cycle in a two-dimensional annulus in transcriptome space suggests that there are essentially two independent groups of genes, the interaction of which drives the cell cycle (*19*). Due to the linearity of our algorithm, the dynamical components DC1 and DC2 represent a core set of genes for these two groups (fig. S8). Together, they comprise 170 genes with significant weights. 52 of these genes are found across all three cell types investigated (HeLa, HEK, 3T3), most of which are well-known to be cell cycle-related. The amount of joint cell cycle genes is in good agreement with other studies comparing multiple cell types (*20, 21*). The well-known cyclin network provides one of the interactions between DC1 and DC2. The representation of the cyclins in DC1 and DC2 is in agreement with their biological function (described in more detail in Methods). Hence, our methods provide a basis for extended mechanistic studies.

The cell’s response to perturbations is described by the stability of the annular manifold – the more stable the manifold is, the faster the cell returns to the unperturbed state. A manifold that is dynamically stable is called an attractor. Mojtahedi et al. (*22*) have shown that the ratio of average gene-to-gene correlation to average cell-to-cell correlation increases with decreasing stability of attractors in transcriptome space. Based on this measure, we found that the stability of the attractor throughout the cell cycle does not change significantly (fig. S9), i.e. the cell types we investigated (HeLa, HEK, 3T3) do not display time points where they are more vulnerable to perturbations. However, we do find consistently that the average pairwise cell-to-cell correlation increases during the cell cycle, reaching its maximum during M phase before a sharp drop during cell division. This is an indicator for a tighter regulation of gene expression of cells in M phase.

Investigating the cloud of data points of single cells, our analysis so far has mapped out the sub-volume of the transcriptome space within which cell cycle dynamics happen. However, this analysis did not yet reveal the shape of the individual trajectories from which these data points are sampled. Within the attractor (the stable manifold), cells might run on a simple circle or follow a more complicated trajectory (i.e. spiraling around the torus) (Fig. 1A). Identifying trajectories requires not only the position of individual cells but also information on the direction of their motion. Since sequencing data contains information about nascent and mature mRNA, transcriptome changes of single cells (their RNA velocity) can be approximately calculated (*23*).

RNA velocity plotted onto the cell cycle reflects the expected order of cell cycle phase clusters and suggests that the attractor is formed by many circles (Fig. 3A). More complicated motion is not supported by the data. The arrows in Fig. 3A represent the direction of motion of individual cells but do not outline a complete trajectory. We can obtain a complete trajectory not for one individual cell but only as a trajectory of an averaged cell (radial averages for each angle bin). If our cell state data and RNA velocity data are consistent, the average RNA velocity should be tangential to the average trajectory. In Fig. 3B, we observe that the average RNA velocities are indeed mostly tangential when plotted onto the average trajectory (also fig. S3D, S4D, S5D, S6D). This strongly suggests that within the two-dimensional projection of the attractor, single cells do in fact move on a simple circle, and that the direction of the motion (but not necessarily the speed) is determined by transcription.

**Fig. 3.**
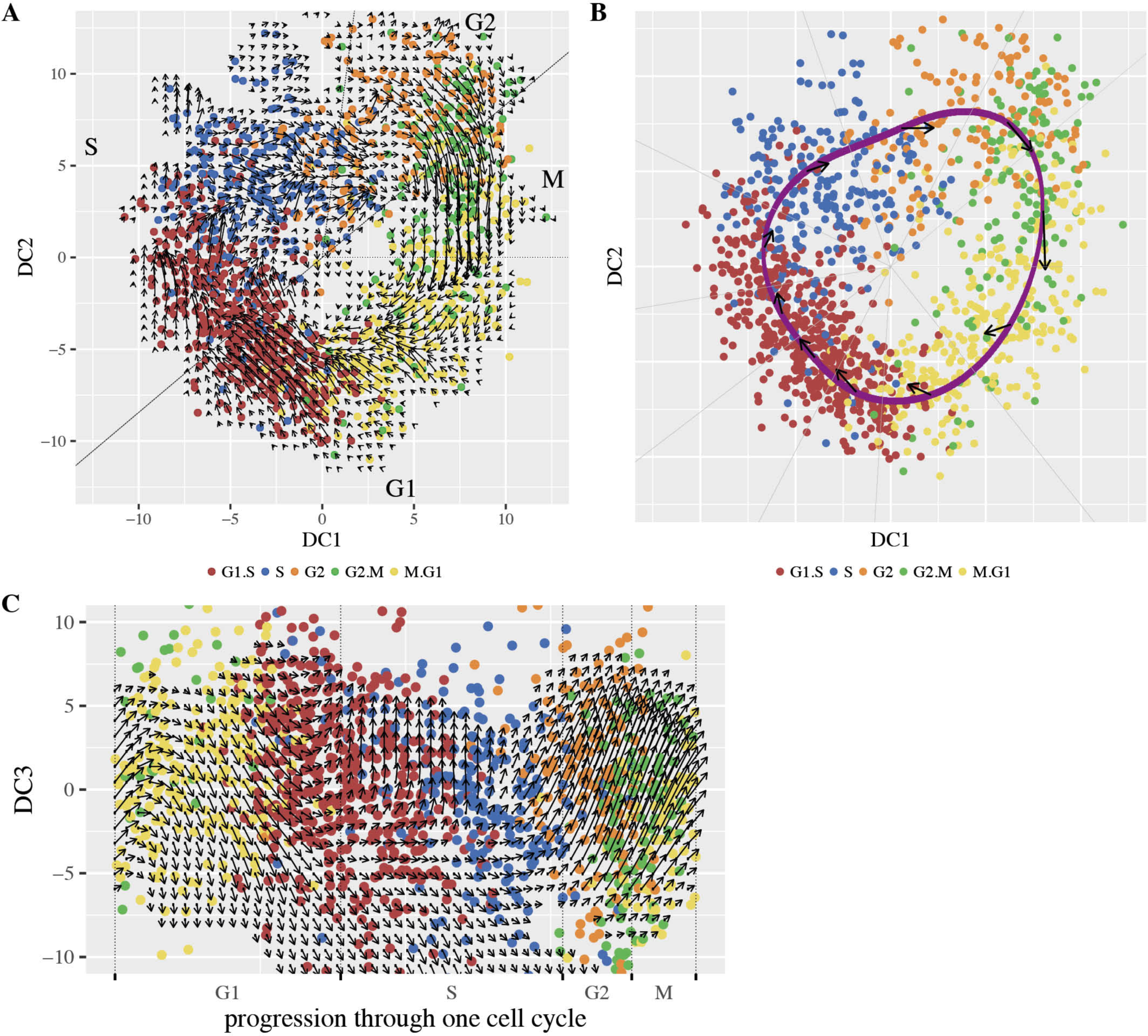
RNA velocity confirms a cyclic motion of the cells and is non-oscillating in the third dimension. (**A**) RNA velocity for each cell is calculated (*23*). We overlay the rotated PCA plot with a 50×50 grid. We assign a weighted average of velocities of surrounding cells with the help of a Gaussian kernel to each grid point (*23*). The estimation of the future state is represented by the arrow head and connected to the originating grid point. Only arrow lengths above a threshold of 0.1 are depicted. (**B**) A spline is placed along the cell cycle approximating an average cell trajectory through the data. We divide the angle into 10 intervals, each containing the same amount of cells (grey borders). For each interval, the average RNA velocity (*23*) in that interval is calculated and plotted onto the average trajectory as a black arrow and shows tangential direction to the average cell cycle. (**C**) RNA velocities in the side of the cylinder (*23*). The cylinder has been cut open at the angular coordinate of the M-G1 transition. The third rotated principal component (DC3) is parallel to the cylinder axis. We do not observe an oscillating motion in the cylinder side. The undulating motion seen in this surface corresponds to a net upward drift after completion of the cell cycle. This indicates that the third dimension does not play a part in the description of the cell cycle. Only arrow lengths above a threshold of 0.4 are shown.

The motion of cells is most coherent during M phase and least directed near the end of S and during G2 phase (Fig. 3A). This points towards a tighter regulation of gene expression during M phase forcing cells through a gene expression tunnel. Cells appear to be more variable in their gene expression when exiting S phase and entering G2 phase. Similarly, we notice more variance of movement direction during the first half of G1 phase and more directed movement once the cells approach the G1-S transition (Fig. 3A). These observations fall in line with the behavior of the coefficient of variation (CV) of the radius along the cell cycle. We find increased CV values during the beginning of G1 phase, the end of S phase and G2 phase (less regulation), and lower CV values during the end of G1 phase and towards M phase (tighter regulation) (fig. S3F, S4F, S5F, S6F, S7).

We also characterized the motion of cells in the direction of the cylinder axis. During G2 and M phase there is a clear upward motion in the direction of the cylinder axis (Fig. 3C). During G1 phase there is a motion in the opposite direction, with a smaller magnitude than the G2 and M phase motion. Hence, each time cells pass through the cycle, they move a little upward but do not return. This motion is a drift parallel to the cylinder axis, which is not periodic and much slower than the motion on the cycle. An inspection of the GO terms associated with the cylinder axis suggests that response to environmental changes (e.g. change of nutrients) and changes of the epigenetic state dominate the processes that cause the motion parallel to the cylinder axis (Table S1). The quality of the separation of cell state dynamics into cell cycle and slower processes depends on the depths of the sequencing data. Data sets with lower depths (i.e. data set 2, data set 3 – see Table S2) do not always show the slow net drift parallel to the cylinder axis but rather some periodic undulations in this direction as well. In those cases, the data sets also exhibit some remaining cell cycle GO terms in the cylinder axis direction, showing that the complete functional isolation of the cell cycle requires sufficient sequencing depth.

In summary, the RNA velocity confirms our assumption that the cell cycle of individual cells is well approximated by a circle and demonstrates that our analysis can separate fast cyclic motion from slow drift, if a sufficient level of detail is achieved in the data. A simultaneous upward motion in the direction of the cylindrical axis transforms the trajectories from circular to helical motion on a hollow cylinder in transcriptome space (Fig. 4).

**Fig. 4.**
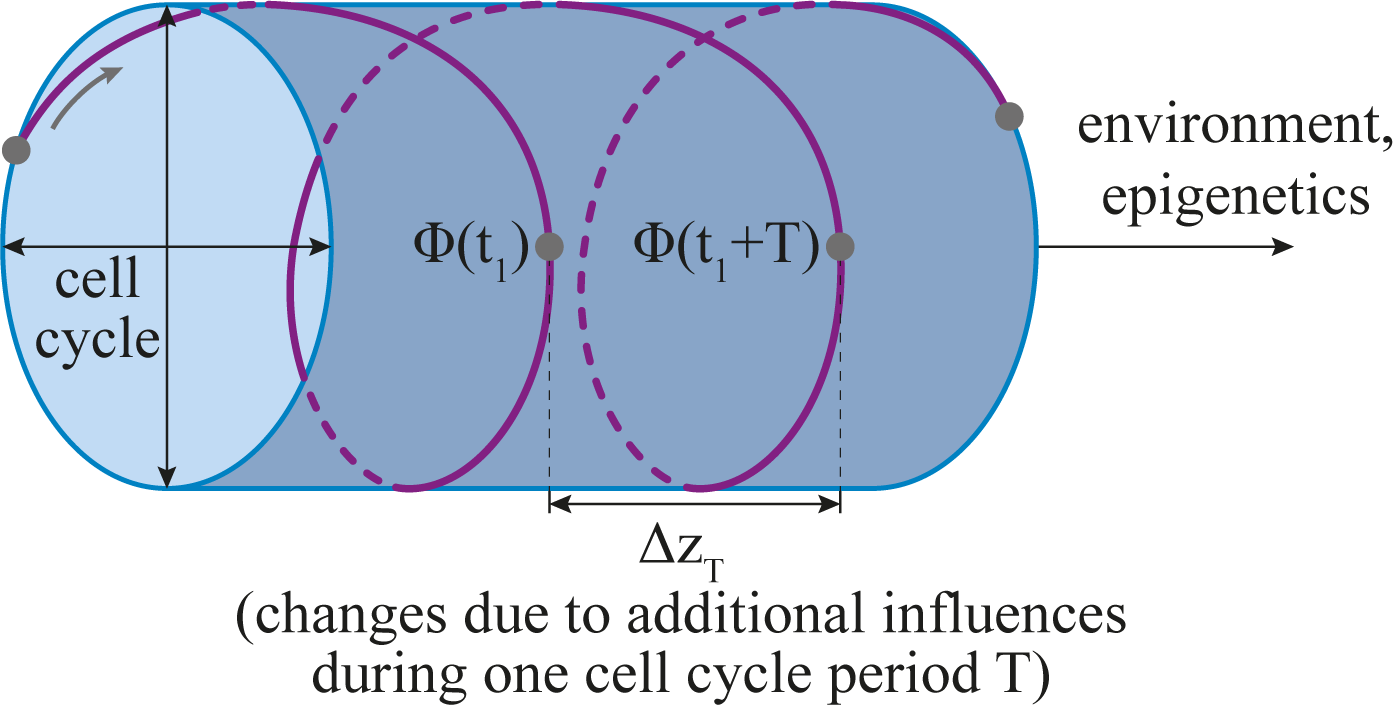
Helical motion of a single cell in transcriptome space. Gene expression changes due to cell cycle can be simplified to a two-dimensional circle by viewing the cylinder from the front or back. Additional cellular processes suggested by GO terms to correspond to epigenetics and environmental changes, cause a helical motion around a hollow cylinder in transcriptome space. During one cell cycle period of time T, the cell will move along the cylinder axis by ∆*z*_*T*_.

Based on the distribution of cells along the cylinder axis, we can divide the three-dimensional cylinder into three parts and analyze the cycle separately in the bottom, middle and top range of the cylinder height. The resulting average trajectories in all three ranges are very similar (fig. S10). We also do not observe clustering of cells with respect to cell cycle phases in the direction of the cylinder axis. Thus, cells appear to be capable of progressing through the cycle independent of influences from cellular processes not represented in DC1 and DC2, such as environmental conditions and epigenetic state, as represented by the direction of the cylinder axis. Due to this isolation of cell cycle functionality into two dimensions and its decoupling from other processes, discounting the cell cycle from data sets via our approach (i.e. subtracting DC1 and DC2) should preserve biological information in the rest of the data accurately.

Our results offer a deep characterization of the transcriptome dynamics of the cell cycle, including a simple method to order unsynchronized cells in time. Our analysis benefits from the relatively high depth of our Drop-seq data. Interestingly, recent data based on sequential single-molecule FISH (which has much higher RNA detection efficiency than single-cell RNA-seq) also supports our findings (*24*). Importantly, our analysis of cell cycle topology is based on analytical methods that are linear and therefore preserve the geometry of the trajectory in gene expression space. These geometrical properties of the trajectory are directly linked to transcriptional regulation. The circular shape of the cell cycle trajectory effectively minimizes curvature. High curvature of the trajectory in transcriptome space would indicate large acceleration of gene expression, achieved by starting or terminating transcription of genes. Such changes generally entail a large regulatory effort by the cell: signaling pathways have to be activated, chromatin rearranged, transcription factors, cofactors and activators recruited, RNA polymerases bound (*7*). The shape of the trajectory shows that the cell cycle has evolved to avoid these efforts for many genes at the same time. Additionally, the simple cycle is the shape of the trajectory guaranteeing that each gene is up- and down regulated not more than once during the cell cycle. Since we obtain this shape in different cell types, this suggests a universal design principle of the cell cycle.

The linearity of our algorithm is in contrast to non-linear analysis and visualization methods (k-nearest-neighbors, UMAP, tSNE), which can be used to flatten more complex manifolds onto a two-dimensional space. It is generally accepted that single-cell transcriptomic profiles characterize an expression manifold embedded in the expression space of all genes. Our work shows that, in our setting, the cell cycle is an independent, two-dimensional manifold within the expression manifold. This begs the question whether the remaining expression manifold can be reduced into further independent submanifolds. Finally, we note that if cells have evolved “optimality principles” to traverse the cell cycle (as we have shown here) it is tempting to speculate that similar optimality principles of gene expression trajectories may have evolved for a large variety of biological systems - in essence for any developmental or cellular differentiation process. Our method and conceptual framework may be useful to discover these principles.

## Acknowledgements

The HeLaS3 AGO2KO monoclonal cell line (*25*) was kindly provided by the laboratory of Valerio Fulci. We would like to thank Christine Kocks for help and advice with FACS strategy and droplet-based sequencing. We thank Seung J. Kim for generating the processed sequencing data, Salah Ayoub for access to and help with additional sequencing experiments and Bastiaan Spanjaard for extensive and helpful comments on the manuscript.

## Authors contributions

NR conceived the research, designed and supervised the experiments. SF carried out the experiments and analysed data. DS designed and carried out the theoretical methods and analysed data. MF designed and supervised the theoretical methods. DS, SF, JPJ, MF, NR wrote the manuscript.

## Data availability

All data is available in the manuscript or the supplementary materials.

## Supplementary Materials

Materials and Methods

Table S1 – S7

Fig S1 – S10

References (25 – 38)

## Materials and Methods

### Conceptual Ideas

#### The transcriptome as a dynamical system

In the context of our considerations, the state of the transcriptome is completely described by the molecule copy number of all species of transcripts in the cell. We can represent the state of the transcriptome of a single cell in a coordinate system with as many axes as there are species of transcripts. The state of the transcriptome is a point in this high-dimensional space. The cell changes its transcriptomic state again and again over time. Hence, among its many other aspects, the transcriptome is also a dynamic system. Change of state is motion along a trajectory in transcriptome space.

We recollect two general results of dynamic systems theory here. Firstly, the trajectory of a dynamical system cannot intersect with itself (*19*). Secondly, the minimum number of dimensions required to embed a trajectory (in conforming with the first point) is a lower bound for the number of ordinary differential equations required to describe the dynamics (*19, 26*), or in other words a lower bound for the number of independent players shaping the trajectory.

Trajectories of a periodic process are closed trajectories. In Fig. 1A, we show examples of such trajectories in a two-dimensional transcriptome space and a three-dimensional space – two resp. three genes participate in these toy dynamics. We consider first the example Fig. 1A.1. Completing it implies regulating gene X up and down once, and the same for gene Y. The next example in Fig. 1A.2 is a cartoon of an extreme case. It requires partial up- and down-regulation of both genes 6 times for completing the cycle – many more up- and down-regulations than the number of participating genes.

The first three-dimensional example (Fig. 1A.3) is more complicated than a circle. It is a type of trajectory found in many dynamical systems and requires at least three dimensions to embed it. The trajectory consists of two loops distinguished mainly by the value of Z. Completing this trajectory once requires more regulation of gene expression than with a circle. It implies regulating gene Z up and down once. Genes X and Y are regulated up and down twice – one time in the ‘lower’ loop and one time in the ‘upper’ loop. Thus, completing this trajectory requires more transcriptional activation and termination than a simple circle.

The example in Fig. 1A.4 is called a torus, and again a common type of trajectory requiring three dimensions to embed it. It also implies more transcription initiations and terminations per cycle than the number of participating genes. The last example (Fig. 1A.5) is an extreme cartoon again, but given the high dimensionality of the transcriptome space, it is a reasonable possibility.

These considerations clearly show there is a relation between the shape of the trajectory in transcriptome space and the regulatory effort - the number of up- and down-regulations of a given gene – required to complete the cycle.

If the trajectory in a high-dimensional space runs on a circle in the side of a cylinder, the trajectory entails only as many up- and down-regulations as there are participating genes. Such a trajectory can also be embedded in a two-dimensional state space. However, the axes do not have the meaning of the number of transcripts of a single gene anymore but describe the amplitude of a group of genes like a principal component or a dynamical component resulting from our analysis (see main text). The genes within one group are regulated in a highly coordinated way but not necessarily synchronously.

We find the cell cycle trajectory in a plane in transcriptome space, i.e. it takes two dimensions to embed it (see main text). This is the minimal number of dimensions required for periodic motion. Hence, essentially two groups of genes interact to drive the cell cycle. The composition of our dynamical components DC1 and DC2 represents a suggestion for these groups (fig. S8). Together, they comprise 170 genes with significant weights for the HeLa data set 1.1 (Table S2), 52 of them are found across all three cell types investigated (HeLa, HEK, 3T3). Positive weights in DC1 correspond to M phase genes, whereas negative weights in DC1 are strongly associated with S phase genes. Simultaneously, genes with positive weights in DC2 are highly correlated to G2 phase, while negative weights are mostly absent in evidence of little cyclic activity at the middle of G1 phase. Consequently, only DC1 contains transcripts for cyclin B (a well-known M phase protein) with positive weights and cyclin E (activated during G1-S transition) in antiphase with negative weight. DC1 also contains transcripts of cyclin A, which is highly expressed during M phase as well as G2 phase. The latter causes it to also have a significant contribution to DC2. The feedbacks between the cyclins mediated by cyclin dependent kinases and other factors represent one of the interactions between DC1 and DC2. Cyclin B1 or B1 and B2 have been shown to be essential for the cell cycle (*27–29*), suggesting that the cyclin network is the only mechanism able to drive cells completely through the cycle. That is in line with the simplicity of the cell cycle trajectory observed in this study.

### Computational Methods

#### Filtering and Cell Cycle Phase Assignment

We use computational methods to identify the phase of the cell cycle a specific cell was in at the moment of measurement. The analysis is based on the principles described in Macosko et al. (*8*) where marker genes for different time points throughout the cell cycle are utilized to assign cells to their cell cycle time point.

We first filter the *m*-by-*n* (genes-by-cells) digital gene expression matrix *S* to ensure every gene is expressed in at least 5 cells and every cell included in the analysis expresses at least 500 genes. We then normalize each column of *S* by the total amount of UMIs *θ*_*j*_ within the *j*-th cell and scale by a factor 10^4^, (*11*). This, we call scaled fraction matrix *SF*:

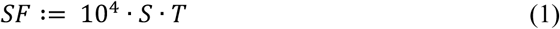

where

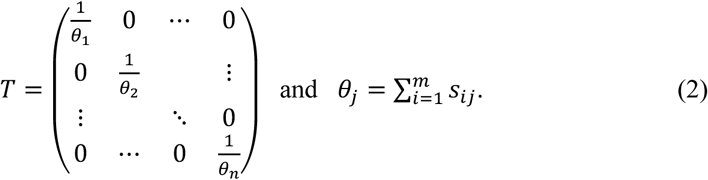

The entries in *SF* are referred to as counts per ten thousand (CPTT). They are displayed in Fig. 2B for specific genes. Next, we take the logarithm of *SF* (*11*). This, we refer to as the logarithmic fraction matrix *LF*:

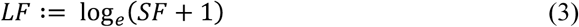

According to Whitfield et al. (*10*), five different cell cycle time points (G1.S, S, G2, G2.M and M.G1) are characterized by specific lists of genes each, which are typically highly expressed at the corresponding cell cycle phase. These are our five marker gene lists, 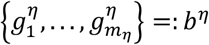 for 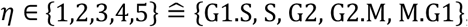, specifying five buckets *b*^*η*^. Genes not appearing in our data set are discarded from this gene list.

The average expression pattern *ξ*_*η*_ of each cell *j* w.r.t. each bucket is defined as the vector whose *j*-th entry is given by

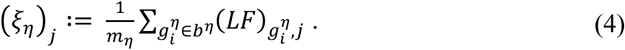

For each row of *LF* that corresponds to a gene 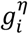 contained in bucket *b*^*η*^, we now calculate

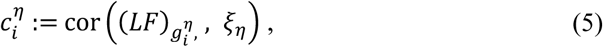

providing information on how well the expression of a single marker gene corresponds to the average expression of its bucket (*8*). We discard all genes 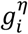 from our buckets for which 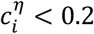 as they are deemed to behave differently than other genes within the bucket and thus do not contribute to inferring cell cycle phases (*8*). This yields the buckets 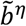 for *η* ∈ {1,2,3,4,5}.

The phase assignment score for cell *j* for phase *η* is given by

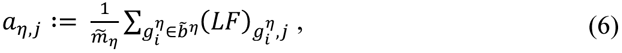

yielding the 5-by-*n* matrix *A* = (*a*_*η,j*_) (*8*). *A* is normalized w.r.t. rows and columns which transforms *A* into a matrix of z-scores (*8*). For each cell *j*, we calculate 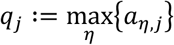 and we declare phase *η*, for which *q*_*j*_ = *a*_*η,j*_ holds, to be the cell cycle phase *η*^*j*^ in which cell *j* is currently located.

We apply two filtering steps. Firstly, we define 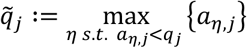 which gives us the second highest phase assignment score for each cell and the associated phase 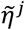. We discard cells for which 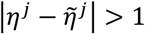 and 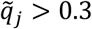. The first condition indicates that the associated phases *η*^*j*^ and 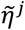 are not neighboring phases while the second condition states that the second highest phase assignment score is significant. These types of cells are suspected doublets as their gene expression peaks in two distant cell cycle phases.

Secondly, we discard cells for which *q*_*j*_ < 0.75 as these cells appear to not exhibit sufficient information for a cell cycle phase assignment.

The data is then again cleaned making sure every gene is expressed in at least 5 cells and every cell expresses at least 500 genes.

In case of data from multiple experiments, the cell cycle phase assignment is done for each experiment individually to avoid dominance of batch effects on the z-scores of the cell cycle phase assignment.

#### Variable Genes

In order to investigate variability within our data set without incorporating information on oscillating genes during the cell cycle from the literature, we obtain variable genes according to the principles from the R package Seurat (*11*):

We calculate the mean expression *ζ*_*i*_ of each gene *i* via

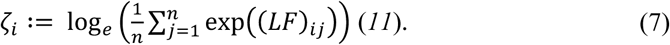

The dispersion *d*_*i*_ of a gene *i* is calculated by taking

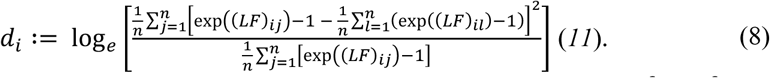

We then compute 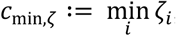, 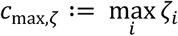 and the step size 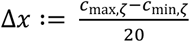. Next, we define 20 buckets 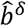 for *δ* ∈ {1, …, 20} such that the *i*-th gene 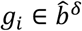 iff

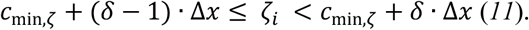

For a specific 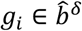, we normalize the dispersion *d*_*i*_ according to all genes within the same bucket 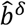:

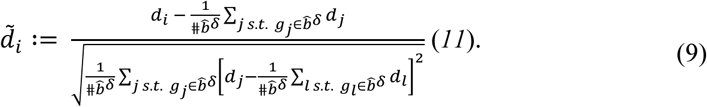

Lastly, we define a gene *i* to be variable iff 0.2 < *ζ*_*i*_ < 4 and 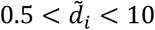. The collection of these genes is denoted by *VG*.

Similar to the cell cycle phase assignment algorithm, we analyze variable genes for each experiment *ds*_*l*_ individually in case the data contains *L* > 1 data sets. There are multiple ways of combining the resulting variable genes of each batch 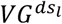 into one set of variable genes *VG*^all^. We have chosen:

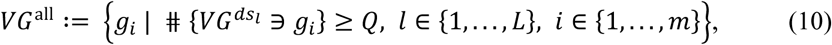

where

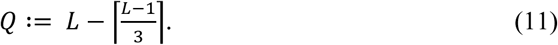

As an example: For *L* = 2 this yields *Q* = 1 and thus

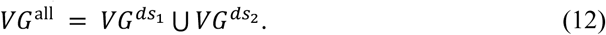

#### Principal Component Analysis

In order to apply PCA, we normalize *LF* (Eq. 3) row-wise so that genes are normalized across all cells. Additionally, we reduce the data set to the variable genes (Eq. 10) giving us the normalized data matrix *N*.

We can write the covariance matrix Cov(*N*^*T*^) of the transposed normalized data matrix *N*^*T*^ as

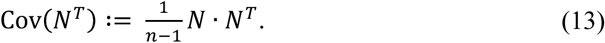

Since Cov(*N*^*T*^) is a real, symmetric, square matrix, we know there exists a matrix *W* such that

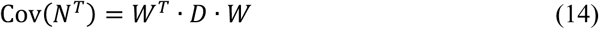

holds, where *D* is a diagonal matrix with the eigenvalues of Cov(*N*^*T*^) as its diagonal elements and where the rows of *W*^*T*^ are the eigenvectors of Cov(*N*^*T*^). *W*^*T*^ is orthogonal and even orthonormal. An entry 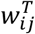 of *W*^*T*^ is called weight *i* for gene *j*.

According to principal component analysis (PCA), we obtain a representation *P* of our data *N* with respect to principal components (PCs) by defining

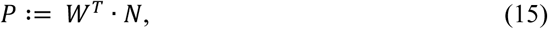

where the rows of *P* are now uncorrelated to one another. A row vector *p*_*k*,_ of *P* contains the PC scores (or amplitudes) of all cells with respect to PC *k* for *k* ∈ {1, …, # *VG*} =: *K*. The pairwise combinations of the first three PC scores for each cell are depicted in (Fig. 1A). The representation *P* of the data *N* according to PCs will be referred to as PC space.

#### Cell Cycle Score

We define a score to judge to what extent a specific PC *k* for *k* ∈ {1, …, ⋕ *VG*} is influenced by the cell cycle. Let *p*_*kj*_ be the *k*-th PC score for cell *j*. We divide all cells *j* for *j* ∈ {1, …, *n*} into five clusters 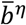, *η* ∈ {1,2,3,4,5} according to their computationally inferred cell cycle phase *η*^*j*^. For a given PC *k*, we then calculate the mean 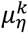 of the PC score for each individual cluster of cells 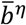:

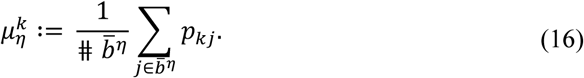

We thus obtain for each PC *k* five different mean values 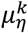.

Our idea is now that in case PC *k* is not influenced by the cell cycle, we should find that these five mean PC scores 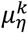 attain similar values since the clustering 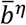 (which is done w.r.t. cell cycle phases) should have a negligible impact on the mean PC scores of the clusters. If on the other hand a PC *k* is influenced by the cell cycle, we expect this to be reflected by differing mean values 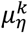. This behavior can be measured by investigating the standard deviation *σ*_*k*_ of the five mean values 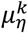

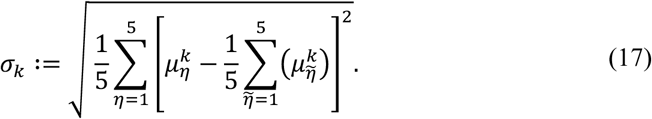

We define *σ*_*k*_ to be our cell cycle score. We note that according to our previous assumptions *σ*_*k*_ is small for a PC *k* that is not influenced by the cell cycle. The least cell cycle influence we would expect for any PC *k* with *σ*_*k*_ = 0. If we assume that the cell cycle does in fact manifest itself within *M* ≪# *VG* PCs, then any PC *k* that is influenced by the cell cycle, should exhibit a significantly higher *σ*_*k*_ than the majority of PCs.

This is a relative score meaning that we are not assigning meaning to the absolute values *σ*_*k*_ of the score. Only if we see significantly higher values in some components than the majority can we hypothesize that these components are influenced by the cell cycle.

#### Rotation of Three-Dimensional Space

We want to rotate the PC space spanned by the first three principal components in order to find a two-dimensional plane that contains the cell cycle. A rotation of three-dimensional space may be executed by a series of two-dimensional rotations with matrices taking the form

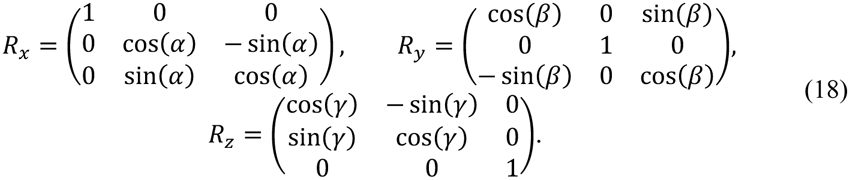

for some angles *α*, *β*, *γ*. The resulting rotation is then given by *R*(*α*, *β*, *γ*) = *R*_*z*_(*γ*) ∙ *R*_*y*_(*β*) ∙ *R*_*x*_(*α*).

Without loss of generality, we can dismiss one of these rotation matrices as all necessary rotations of the space can be achieved by a combination of two angles. We choose *γ* = 0, yielding *R*_*z*_ = *I*_3_.

More generally, we can rotate a three-dimensional subspace of a larger vector space (with dimension>3) by filling out all other dimensions not involved in the rotation by the identity matrix. An example of a ration matrix *R* of the three-dimensional subspace spanned by dimension 1, 4, 6 in a 6-dimensional space:

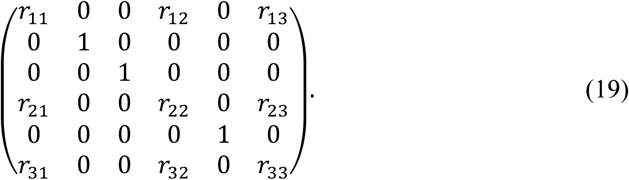

Our goal is to find appropriate angles *α*, *β* such that the direction vector *ω* of the axis of the cylinder forming the manifold is in the direction of the third rotated component

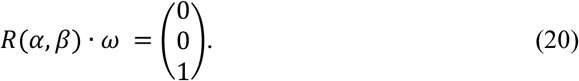

Given *ν*, the entries of the rotation matrices system can be determined like

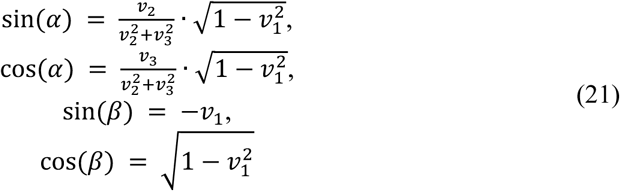

#### Finding the Optimal Rotation

We want to rotate the PC space in an unsupervised manner. Our optimization is that after rotation of a three-dimensional subspace spanned by components *k*_1_, *k*_2_, *k*_3_ for *k*_*i*_ ∈ {1, …, # *VG*} the cell cycle score 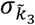 (Eq. 17) is minimal in the new third component 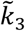. This condition derives from considering a triplet of mean PC scores 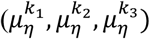 (Eq. 16) as a point *γ*_*η*_ within our three-dimensional subspace for *η* ∈ {1,2,3,4,5}. Each of the five cell cycle time points (G1.S, S, G2, G2.M and M.G1) yields one such point *γ*_*η*_. We now attempt to place all of these five points *γ*_*η*_ into a single plane. Minimizing the distance of the five points *γ*_*η*_ to that plane is equivalent to minimizing the cell cycle score for the vector orthogonal to the plane. The fact that such a plane exists is non-trivial. We will refer to the orthogonal vector corresponding to the plane as the viewing axis.

The pair (*α*,*β*) defines a solid angle. We do a two-step optimization. First, we divide the total solid angle of 4*π* into 10.000 bins of equal size. Utilizing the golden spiral algorithm (also referred to as spherical Fibonacci grid) (*30, 31*), we generate 10.000 approximately equidistantly spaced points on a unit sphere. Each of these points is a potential viewing axis. For each of them we calculate the corresponding cell cycle score. The axis *ω*_*i*_, *i* ∈ {1, …, 10.000} associated with the lowest cell cycle score is chosen to be optimal.

As a second step, we refine the grid of potential viewing axes in a small neighborhood of *ω*_*i*_ by roughly the factor 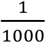. Again, we find the viewing axis 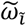 associated with the lowest cell cycle score and choose this 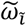 to be the viewing axis that becomes the vector 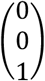 after rotation of the three-dimensional subspace.

#### Generalization to Sequence of Rotations and Selection of Significant CC Components

So far, we have always assumed that PCA manages to place the cell cycle drivers within the first three dimensions. This is unfortunately only true for sufficiently deep sequenced data sets. We have investigated multiple Drop-seq data sets from HeLa, HEK and 3T3 cells where we find significant cell cycle scores for more than three PCs. Therefore, it is necessary that we advance from a single rotation of a three-dimensional subspace to a sequence of three-dimensional rotations. We note that combining two rotation matrices *R*_1_, *R*_2_ again yields a rotation matrix *R* = *R*_2_ ⋅ *R*_1_.

We consider the cell cycle score for the first 50 principal components. We need to judge which of these components are significantly influenced by the cell cycle. This comes down to an outlier detection problem. We would in general expect to obtain most cell cycle scores close to zero with only a handful significantly higher scores, implying cell cycle influence for those few components. We deviate from the usual way of outlier detection via sample mean/sample standard deviation as our sample standard deviation of only 50 values would be heavily influenced by strong outliers. Instead, for the vector 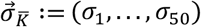 (see Eq. 17), 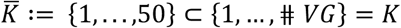, we consider the median absolute deviation (MAD)

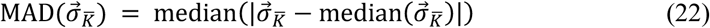

and define 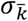 for 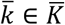 to be an outlier iff

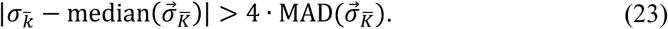

Let 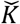 be the collection of 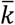 for which 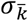 is an outlier. Then any PC 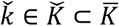 is considered to be a PC on which the cell cycle has significant influence.

Our goal is to place the cell cycle influence into the first two components. Therefore, the first two components always span the first two dimensions of the three-dimensional subspace we rotate. The third dimension is spanned by a PC 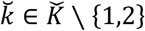.

As an example, we assume our outlier detection found that PC1, PC2, PC3, PC5 and PC8 have significant cell cycle scores. This yields 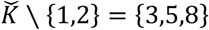 and implies that we require three subsequent three-dimensional rotations. The first step is the same as described previously: We select the three-dimensional space spanned by PC1, PC2 and PC3, we find the optimal viewing axis for this subspace and rotate the data set accordingly by a matrix *R*_1_. This yields rotated-PC1, rotated-PC2 and rotated-PC3 where the cell cycle score of rotated-PC3 was minimized and the cell cycle effects exhibited by PC3 previously were ideally included into rotated-PC1 and rotated-PC2. In the next step, we select the three-dimensional subspace spanned by rotated-PC1, rotated-PC2 and PC5 and find the optimal viewing axis such that the cell cycle score is minimal in rotated-PC5. We obtain *R*_2_. Finally, this is repeated with the newly rotated-PC1, newly rotated-PC2 and PC8 yielding *R*_3_. In total, we have a sequence of three three-dimensional rotations *R*_1_, *R*_2_, *R*_3_ which when combined are in fact realized by a single rotation matrix *R* = *R*_3_ ⋅ *R*_2_ ⋅ *R*_1_.

We find that with this method, we are able to isolate cell cycle effects into just two dimensions for all data sets investigated (fig. S1B, S1C, S3B, S4B, S5B, S6B). The algorithm is not influenced by batch effects and will ignore such effects as long as the relevant cell cycle information is present and contained within the first 50 PCs. We have set the boundary of 50 PCs as we have not yet found any data set that had significant cell cycle scores past the 50th PC. The algorithm can be extended to include as many PCs as desired. Only the detection of outliers has to then be adjusted to account for additional data points influencing the outlier detection algorithm.

In the end, we find a new representation *P*_*R*_ of the data *N* by multiplying a rotation matrix *R* from the left onto the representation *P* (Eq. 15) via

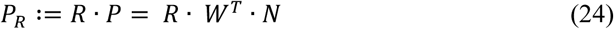

where *R* is a sparse orthogonal matrix which causes cell cycle effects to be maximized in the first two dimensions. The representation *P*_*R*_ of the data *N* according to rotated PCs will be referred to as rotated PC space.

#### Dynamic components

The dynamics of the cell cycle is the dynamics of the mRNA and protein concentrations of the cell. We restrict our analysis to the mRNA concentrations. Neglecting noise, it can be described by a large system of ordinary differential equations

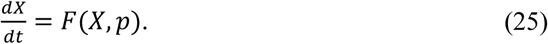

Here *X* denotes the vector of the mRNA concentrations, *p* a vector of parameter values and *t* denotes the time variable. The dependence on *p* captures also cell variability. In general, the time course of *X* on the manifold can be described by a system of differential equations for abstract variables *A* = (*a*_1_, …, *a*_*M*_) with fewer components *a*_*i*_ than the large number *m* of mRNAs:

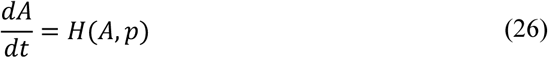

The original data are related to the abstract variables by algebraic functions *G*

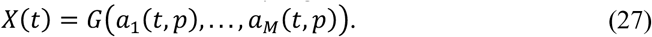

Such a description is useful, if very few *a*_*i*_ provide a good approximation of the time course, i.e. *M* ≪ *m*. *M* is an upper limit for the dimension of the manifold. There is a variety of methods of finding the abstract variables (*26, 32*). Our results show, that the cell cycle dynamics (motion on the manifold) can be represented in good approximation with *M* = 2, described by differential equations for *a*_1_ and *a*_2_:

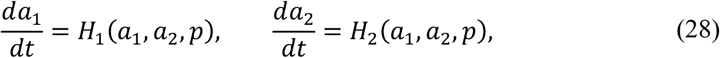

and a particularly simple function *G*:

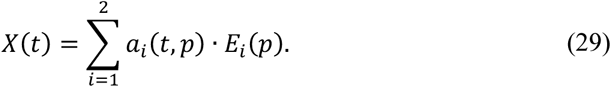

*E*_1_ and *E*_2_ are sums of principal components and are called dynamical components (DCs). The rotated PC1 and rotated PC2 are one of several possible choices of *E*_1_ and *E*_2_. We therefore denote rotated PC1 and rotated PC2 by DC1 and DC2, respectively. Other possible choices follow from rotated PC1 and rotated PC2 by rotation around the cylinder axis (rotated PC3 direction).

#### Synchronizing Cell Cycle to Cell Division

In order to compare different data sets, we want to find a way to synchronize the obtained cell cycle to a known time point that exists in all data sets. Cell division is present in all cell types we are investigating. Furthermore, we can approximate the moment of cell division by investigating the amount of mRNA transcripts contained within cells along the cell cycle which makes cell division an ideal candidate to align our data sets to.

More specifically, we consider the cell cycle displayed by DC1 and DC2 and divide the data into 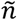 bins containing 30 cells each. For each bin, the average total UMI count is calculated providing us with a time course 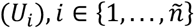 of average total UMI counts along the cell cycle. We take the minimum and maximum of (*U*_*i*_) and construct a linear function *h*(*x*) such that 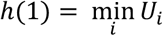 and 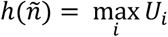. We now consider all permutations *π*_*l*_, 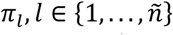 of the set 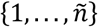 which periodically shift every element by *l* − 1. Then we search for the permutation *π*_*l*_ which attains the minimum in squared residuals between the time course and the linear function:

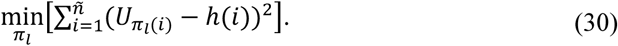

This is a simple but efficient approximation for the bin *l* at the start of which the cell division is most likely to take place due to the fact that we expect a sudden drop in average total UMI counts between cells about to divide and the ones that just divided. The increase in average UMI counts per cell is in reality not linear but we have seen during analysis of multiple data sets that this approximation is sufficient.

Finally, let *α*_*x*_ be the minimal angle a cell attains in polar coordinates within bin *l* and let *α*_*x*−1_ be the maximum angle of all cells from bin *l* − 1. We rotate the two-dimensional cell cycle about an angle 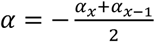, thereby placing the time point of cell division onto the positive axis of DC1.

#### Phase Space Density and Speed Along Trajectories

Phase space density is a measure for how many data points are clustered around a certain area within the phase space. Considering the cell cycle given by DC1 and DC2 in polar coordinates (*ϕ*, *r*), we can estimate the radius *r*_*c*_(*ϕ*) of an idealized cyclic cell through the center of mass along the cell cycle. While this immediately provides us with a good estimate of the shape of the trajectory of an individual cell in phase space, it does not tell us what the function *ϕ*(*t*), the angle *ϕ* depending on time *t*, looks like.

We hypothesize that due to the large amount of data points within our data, we should have sufficient and continuous coverage of the cell cycle. Due to the fact that experiments were done on asynchronous data, we assume that the distribution of data points along the cell cycle is uniform in time across the cell cycle period. We divide the cell cycle period into equal bins of size ∆*t*. The angle ∆*ϕ* a cell moves during the time ∆*t* is proportional to the angular velocity *ν*: ∆*ϕ* = *ν*∆*t*. We can conclude that the velocity 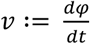, with which we progress along a curve such as *r*_*c*_(*ϕ*), is inversely proportional to phase space density, since ∆*t* is proportional to the number of cells within a time bin. Hence, we progress more slowly along a given trajectory where there is a large amount of data points present and vice versa.

#### Incorporating Cell Cycle Phase Durations From Literature Into Phase Space Plots

There are multiple publications on measuring the lengths *l* of cell cycle phases. For HeLa and 3T3 cell lines, we obtain values for cell cycle phase lengths from Hahn et al. (*33*) and for HEK293 data sets from Cheng et al. (*34*) These cell cycle phase lengths and their location in our plots are to be understood as rough estimates. We note that we specifically do not observe discrete switches from one phase to another but rather continuous transitions between them. The notations of cell cycle phases were created by scientists in order to group processes and facilitate description of such.

Since we have previously defined the time point of cell division within our data, we can equate this time point to the transition from M to G1 phase. In the previous section, we have argued that we can relate information from literature about time durations to information taken from our cell cycle in phase space by considering the density of data points along the cell cycle in transcriptome space. We order the cells according to their angle in polar coordinates. From the first cell after cell division, we define the transition G1-S to take place after *x* cells where *x* can be computed from

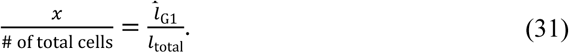

The mean angle of the cells *x* and *x* + 1 gives us an estimate of where in our phase space the transition G1-S takes place. We repeat this step for all remaining transitions analogously.

#### RNA Velocity Analysis

La Manno et al. (*23*) introduced RNA velocity as a concept of distinguishing between unspliced and spliced RNA in order to extrapolate cell states to a future time point. We implement the approximation model I from the supplement of La Manno et al. (*23*) into our analysis. Let *S*_*ij*_(*t*) be the number of spliced transcripts of a specific gene *i* present in a cell *j* dependent on time *t*. The assumption under model I is that the time derivative of *s*_*ij*_ is constant

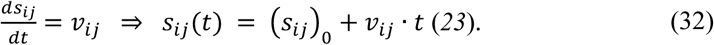

The velocity matrix *V* is estimated as is shown in the supplement of La Manno et al. (*23*). We set 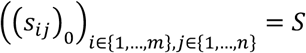 where *S* is our raw data matrix (see Methods "Filtering and Cell Cycle Phase Assignment"). We choose *t* = 1 and we obtain the extrapolated state matrix *S*_ex_ via

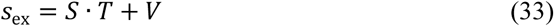

(see Eqs. 2, 32). Next, we transform the data to *LF*_ex_ (Eq. 3) similarly as before

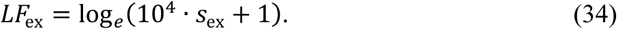

We then normalize *LF*_ex_ for each gene across all cells according to the mean and standard deviation of each gene in *LF*_ex_ and limit ourselves to the variable genes found during our previous analysis which yields the normalized extrapolated data matrix *N*_ex_. Lastly, we transform the data points into the rotated PC space where the first two dimensions represent the cell cycle:

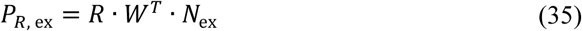

(Eqs. 15, 24).

Due to the high noise level in the unspliced data, we have to incorporate a smoothing grid on top of the DC1-DC2-plot in order to obtain relevant information about the direction of motion of the cells. The calculation of the grid is again done as described in the supplement of La Manno et al. (*23*), incorporating a Gaussian kernel function

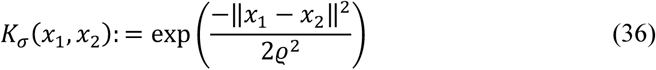

and defining the displacement of a grid point 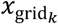 via

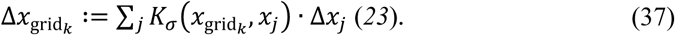

The smoothing parameter *ϱ* (Eq. 36) can be chosen freely. We take care to find a balance between smoothing enough to get a reasonable idea of the general motion of the system but at the same time taking care not to eliminate too much noise so that there is room for interpretation of the strength of cyclic motion at different time points during the cell cycle. In Fig. 3A, we have chosen *ϱ* = 0.6 (Eq. 36).

#### RNA Velocity in the side of the cylinder

In order to show that we isolated cell cycle into just two dimensions, we investigate also the motion of the cells parallel to the cylinder axis. A cylinder can be described by the angle *ϕ* and radius *r* of its base and the height corresponding to the direction of DC3 in our representation of the data..

State changes in *ϕ*-direction are calculated by calculating polar coordinates for our data and the extrapolated data in the DC1-DC2-plot. The changes in height-direction are given by the changes in DC3 between the data and the extrapolated state. Due to high noise levels, we again apply a grid smoothing as outlined before (*23*). The only difference is that we apply a scaling factor to *ϕ* in order to make the Gaussian kernel approximately symmetrical. The scaling factor we choose is the mean value of the radius of all data points in the DC1-DC2-plot. The displacement value for grid points can then be scaled back by the same factor so that we can display the results in the *ϕ*-DC3-plane as shown in Fig. 2B. Here we choose *ϱ* = 2 (Eq. 36).

#### Stability Index of the Attractor

If we compare the attractor (a stable manifold) to a water slide, then a person going down the water slide corresponds to a cell going through the cell cycle. The cell runs along the main path at the bottom of the channel, but it also veers towards the sides. The steepness of the walls, or the strength with which cell is pushed back towards the middle, is called the attractor stability. The steeper the walls, the faster perturbations decay and the faster cells return to the attractor, hence the more stable it is.

The index for critical state transitions *I*_*C*_ introduced in Mojtahedi et al. (*22*) is defined via

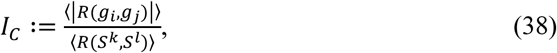

where *g*_*i*_ are gene vectors, *S*^*k*^ are cell states and 〈*R*(…, …)〉 is the average of all pairwise correlations (we utilize Spearman correlation in our analysis) (*22*). We only include pairwise correlation values with a *p*-value smaller than 0.05.

In polar coordinates, we divide the two-dimensional cell cycle plot with respect to angle into 10 bins which contain the same number of cells each. For each of these bins we calculate the stability index *I*_*c*_ as introduced by Mojtahedi et al. (*22*).

The stability index increases before a critical state transition due to the fact that gene-gene correlation occurs coordinately and therefore increases on average, while the average cell-cell correlation decreases because cells are more variable during transitions than in steady state. Both of these effects would cause an increase in *I*_*c*_ making the stability index an appropriate measure for state transitions.

Our analysis of the behavior of *I*_*c*_ throughout the cell cycle implies that there is no critical state transition detected as the progression observed in fig. S9C is homogeneous. Small changes are attributed to noise rather than orchestrated behavior (fig. S9C). We do however observe consistent increase in average cell-cell correlation throughout the cell cycle (fig. S9B). We reach the highest values when entering M phase and observe a sharp drop following cell division. This coincides with other observations mentioned in the main text and supports the claim that cells are more tightly regulated during M phase where they are pushed through a gene expression tunnel whereas they are more variable just after cell division.

#### GO Term Analysis

Since the additional rotation *R* (Eq. 24), which we apply after PCA, is a linear algorithm, the weights *W*^*T*^ (Eq. 15) generated by PCA can be transformed linearly as well by considering the rotated weights 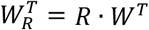. We analyze the rotated weights from *W*_Ò_ and the difference to original weights from *W* in order to gain an insight into the cyclic object we found in two dimensions.

A GO term analysis (*16, 17*) of the first three rows of *W*, which are the weights generating the first three PCs, shows that all three PCs are highly dominated by cell cycle processes (see Table S3, S4, S5). We see highly significant *p*-values suggesting that all three PCs are vital for the description of the cell cycle.

On the other hand, for the weights in *W*_*R*_ we observe that while DC1 and DC2 are still dominated by cell cycle effects with highly significant *p*-values, DC3 is completely free of cell cycle GO terms (see Table S1, S6, S7) (*16, 17*).

Furthermore, we have investigated additional rotated PCs of lower order and have not found any GO terms (*16, 17*) related to the cell cycle with a p-value < 10^−5^. This is another strong indication that we have succeeded in isolating cell cycle effects into only two dimensions.

### Experimental Data

#### Single-cell sequencing for data set 1: Drop-seq procedure, single-cell library generation and sequencing

The Drop-seq runs and library preparation were performed as described in Alles et al. (*9*) on a self-built Drop-seq set up (*8*).

HeLaS3 cells and HeLaS3 AGO2KO cells were grown to the logarithmic phase, pelleted by centrifugation (300 g, 5 minutes), fixed with 80% cold methanol while mildly vortexing and kept on ice until the run. Fixed cells were prepared for the Drop-seq run by centrifugation (1000 g, 5 min) and resuspension in 1 ml of PBS-BSA 0.01% + RiboLock (ThermoFisher) (0.8 U/µl), followed by another centrifugation (1000 g, 5 min) and resuspension in 0.5 ml of PBS-BSA 0.01% + RiboLock (ThermoFisher) (0.8 U/µl). Then, cells were passed through a cell strainer (35 µm), counted, diluted with PBS-BSA 0.01% to a concentration of 100 cells/µl and transferred into a syringe to be loaded on the Drop-seq apparatus. After mixing with lysis buffer, this corresponds to a final concentration of 50 cells/µl in the droplets.

The two single-cell libraries from the HeLaS3 and HeLaS3 AGO2KO cells (1.8 pM, final insert sizes 700 bp) were sequenced in paired end mode on Illumina Nextseq500, together with two other libraries, yielding ~49*10^6^ read pairs for the HeLaS3 library and ~45*10^6^ read pairs for the HeLa AGO2KO library.

A second sequencing run was performed with two multiplexed libraries prepared from a subpool of cells (50%) in order to obtain deeper sequencing data from less cells (1.8 pM, final insert sizes 645 and 628 bp). This yielded ~168*10^6^ read pairs for the HeLaS3 library and ~186*10^6^ read pairs for the HeLa AGO2KO library.

Read 1: 20 bp (bases 1-12 cell barcode, bases 13-20 UMI; Drop-seq custom primer 1 “Read1CustSeqB”), index read: 8 bp, read 2 (paired end): 64 bp).

#### Single-cell sequencing for data set 2: Drop-seq procedure, single-cell library generation and sequencing

The Drop-seq runs and library preparation were performed as described in Alles et al. (*9*) on a self-built Drop-seq set up (*8*).

HeLaS3 cells were grown to the logarithmic phase, pelleted by centrifugation (300 g, 5 minutes), fixed with 80% cold methanol while mildly vortexing and kept on ice until the run. Fixed cells were prepared for the Drop-seq run by centrifugation (1000 g, 5 min) and resuspension in 1 ml of PBS-BSA 0.01% + RiboLock (ThermoFisher) (0.8 U/µl), followed by another centrifugation (1000 g, 5 min) and resuspension in 0.5 ml of PBS-BSA 0.01% + RiboLock (ThermoFisher) (0.8 U/µl). Then, cells were passed through a cell strainer (35 µm), counted, diluted with PBS-BSA 0.01% to a concentration of 100 cells/µl and transferred into a syringe to be loaded on the Drop-seq apparatus. After mixing with lysis buffer, this corresponds to a final concentration of 50 cells/µl in the droplets.

The single-cell final library (1.8 pM, final insert sizes 532bp) was sequenced in paired end mode on Illumina Nextseq500 75 cycles high output, together with three other libraries, yielding ~100*10^6^ read pairs for the HeLaS3 library.

Read 1: 20 bp (bases 1-12 cell barcode, bases 13-20 UMI; Drop-seq custom primer 1 “Read1CustSeqB”), index read: 8 bp, read 2 (paired end): 64 bp).

#### Processing of Raw Sequencing Data Sets

The sequencing quality was assessed by FastQC v.0.11.2 (*35*). We used the Drop-seq tools v.2.0.0 (*8*) to tag the sequences with their corresponding cell and molecular barcodes, to trim poly(A) stretches and potential SMART adapter contaminants and to filter out barcodes with low-quality bases. The reads were then aligned to a GRCh38 reference genome (*36*), using STAR v.2.6.0c (*37*) with default parameters and sorted using samtools v.1.9 (*38*).

The Drop-seq tool was further exploited to add gene annotation tags to the aligned reads and to identify and correct some of the bead synthesis errors. The number of cells was determined by extracting the number of reads per cell, then plotting the cumulative distribution of reads against the cell barcodes ordered by descending number of reads and selecting the inflection knee point of the distribution using dropbead v.0.25 (*9*). Finally, the DigitalExpression tool (*8*) was used to obtain the digital gene expression (DGE) matrix for each sample. DGE matrix with only intronic reads was created by specifying the list of functional annotations.

---

**Table S1.** First 20 entries obtained by GO term analysis (*16, 17*) of the weights associated with DC3 are completely free of any cell cycle terms. None of the significant GO terms have any direct association with the cell cycle, indicating that we in fact managed to eliminate any cell cycle influence in the third dimension, which represents the cylinder axis from Fig. 1A.

| GO term | Description | P-value | FDR q-value | Enrichment (N, B, n, b) |
| --- | --- | --- | --- | --- |
| GO:0006457 | protein folding | 1.51E-8 | 1.23E-4 | 7.36 (1611,35,75,12) |
| GO:0010033 | response to organic substance | 2.25E-8 | 9.14E-5 | 2.77 (1611,233,75,30) |
| GO:0006810 | transport | 3.01E-8 | 8.16E-5 | 2.15 (1611,410,75,41) |
| GO:0051234 | establishment of localization | 1.55E-7 | 3.16E-4 | 2.04 (1611,432,75,41) |
| GO:0042221 | response to chemical | 1.94E-7 | 3.15E-4 | 2.41 (1611,285,75,32) |
| GO:0032940 | secretion by cell | 2.88E-7 | 3.91E-4 | 3.61 (1611,113,75,19) |
| GO:0051179 | localization | 5.12E-7 | 5.96E-4 | 1.87 (1611,505,75,44) |
| GO:0006887 | exocytosis | 7.56E-7 | 7.69E-4 | 3.76 (1611,97,75,17) |
| GO:0046903 | secretion | 7.77E-7 | 7.03E-4 | 3.40 (1611,120,75,19) |
| GO:0065008 | regulation of biological quality | 1.06E-6 | 8.61E-4 | 2.16 (1611,338,75,34) |
| GO:0034975 | protein folding in endoplasmic reticulum | 1.11E-6 | 8.22E-4 | 17.90 (1611,6,75,5) |
| GO:0002376 | immune system process | 1.11E-6 | 7.54E-4 | 2.60 (1611,215,75,26) |
| GO:0045055 | regulated exocytosis | 1.63E-6 | 1.02E-3 | 3.78 (1611,91,75,16) |
| GO:0002697 | regulation of immune effector process | 2.37E-6 | 1.38E-3 | 5.37 (1611,44,75,11) |
| GO:0050896 | response to stimulus | 3.05E-6 | 1.66E-3 | 1.79 (1611,515,75,43) |
| GO:0050778 | positive regulation of immune response | 4.61E-6 | 2.35E-3 | 4.60 (1611,56,75,12) |
| GO:0002684 | positive regulation of immune system process | 5.4E-6 | 2.59E-3 | 3.91 (1611,77,75,14) |
| GO:0070972 | protein localization to endoplasmic reticulum | 6.18E-6 | 2.79E-3 | 10.74 (1611,12,75,6) |
| GO:0016192 | vesicle-mediated transport | 6.25E-6 | 2.68E-3 | 2.51 (1611,205,75,24) |
| GO:0002274 | myeloid leukocyte activation | 6.34E-6 | 2.58E-3 | 3.86 (1611,78,75,14) |

**Table S2.** Summary of sequencing data analyzed.

| ID | cell type | availability | #cells | variable genes utilized? | #genes used during PCA | mean UMI count | displayed in figure/table |
| --- | --- | --- | --- | --- | --- | --- | --- |
| data set 1.1 | HeLaS3 | https://www.ncbi.nlm.nih.gov/geo/query/acc.cgi?acc=GSE142277<br>records private; private review token will be generated upon request | 1.348 | Yes | 1.536 | 14.824 | Fig. 1-3, Table S1,S3-S7, fig. S1,S7,S9,S10, **S8** |
| data set 2 | HeLaS3 | https://www.ncbi.nlm.nih.gov/geo/query/acc.cgi?acc=GSE142356<br>records private; private review token will be generated upon request | 2099 | Yes | 938 | 6146 | fig. S3,S9,S10 |
| data set 3 | HEK293T | Alles et al., BMC Biology (9) | 754 | Yes | 1.545 | 9.515 | fig. S4,S9,S10 |
| data set 4 | 3T3 | Alles et al., BMC Biology (9) | 741 | Yes | 1.306 | 10.018 | fig. S5,S9,S10 |
| data set 1.2 | HeLaS3 | https://www.ncbi.nlm.nih.gov/geo/query/acc.cgi?acc=GSE142277<br>records private; private review token will be generated upon request | 1.348 | No | 10.720 | 14.824 | fig. S6,S9 |

**Table S3.** First 20 entries obtained by GO term analysis (*16, 17*) of the weights associated with PC1 are dominated by the cell cycle.

| GO term | Description | P-value | FDR q-value | Enrichment (N, B, n, b) |
| --- | --- | --- | --- | --- |
| GO:0051301 | cell division | 8.4E-33 | 6.84E-29 | 8.66 (1611,93,82,41) |
| GO:0022402 | cell cycle process | 2.69E-32 | 1.1E-28 | 5.19 (1611,208,82,55) |
| GO:1903047 | mitotic cell cycle process | 5.32E-29 | 1.44E-25 | 5.84 (1611,158,82,47) |
| GO:0007088 | regulation of mitotic nuclear division | 8.85E-23 | 1.8E-19 | 10.42 (1611,49,82,26) |
| GO:0051783 | regulation of nuclear division | 6.77E-22 | 1.1E-18 | 9.82 (1611,52,82,26) |
| GO:0010564 | regulation of cell cycle process | 4.2E-21 | 5.7E-18 | 5.01 (1611,157,82,40) |
| GO:0051726 | regulation of cell cycle | 2.36E-20 | 2.74E-17 | 4.24 (1611,204,82,44) |
| GO:0007346 | regulation of mitotic cell cycle | 8.96E-19 | 9.11E-16 | 5.21 (1611,132,82,35) |
| GO:0051983 | regulation of chromosome segregation | 3.08E-18 | 2.78E-15 | 10.31 (1611,40,82,21) |
| GO:0007017 | microtubule-based process | 1.43E-16 | 1.16E-13 | 5.25 (1611,116,82,31) |
| GO:0007059 | chromosome segregation | 1.36E-15 | 1.00E-12 | 11.13 (1611,30,82,17) |
| GO:0098813 | nuclear chromosome segregation | 2.43E-15 | 1.65E-12 | 14.48 (1611,19,82,14) |
| GO:0000226 | microtubule cytoskeleton organization | 3.42E-15 | 2.14E-12 | 6.22 (1611,79,82,25) |
| GO:0051276 | chromosome organization | 4.06E-15 | 2.36E-12 | 5.87 (1611,87,82,26) |
| GO:1902099 | regulation of metaphase/anaphase transition of cell cycle | 9.91E-15 | 5.38E-12 | 12.28 (1611,24,82,15) |
| GO:0030071 | regulation of mitotic metaphase/anaphase transition | 9.91E-15 | 5.04E-12 | 12.28 (1611,24,82,15) |
| GO:0010965 | regulation of mitotic sister chromatid separation | 2.38E-14 | 1.14E-11 | 11.79 (1611,25,82,15) |
| GO:1905818 | regulation of chromosome separation | 2.38E-14 | 1.08E-11 | 11.79 (1611,25,82,15) |
| GO:1901987 | regulation of cell cycle phase transition | 1.73E-13 | 7.4E-11 | 5.11 (1611,100,82,26) |
| GO:0006996 | organelle organization | 2.86E-13 | 1.16E-10 | 2.74 (1611,330,82,46) |

**Table S4.** First 20 entries obtained by GO term analysis (*16, 17*) of the weights associated with PC2 are similarly dominated by the cell cycle.

| GO term | Description | P-value | FDR q-value | Enrichment (N, B, n, b) |
| --- | --- | --- | --- | --- |
| GO:0022402 | cell cycle process | 2.23E-25 | 1.82E-21 | 4.49 (1611,208,88,51) |
| GO:1903047 | mitotic cell cycle process | 7.83E-25 | 3.19E-21 | 5.21 (1611,158,88,45) |
| GO:0051726 | regulation of cell cycle | 7.91E-18 | 2.15E-14 | 3.86 (1611,204,88,43) |
| GO:0010564 | regulation of cell cycle process | 9.28E-17 | 1.89E-13 | 4.31 (1611,157,88,37) |
| GO:0007346 | regulation of mitotic cell cycle | 1.37E-16 | 2.23E-13 | 4.72 (1611,132,88,34) |
| GO:0051301 | cell division | 1.6E-16 | 2.17E-13 | 5.71 (1611,93,88,29) |
| GO:0051783 | regulation of nuclear division | 1.38E-14 | 1.61E-11 | 7.39 (1611,52,88,21) |
| GO:0007088 | regulation of mitotic nuclear division | 5.1E-14 | 5.19E-11 | 7.47 (1611,49,88,20) |
| GO:1901987 | regulation of cell cycle phase transition | 1.33E-13 | 1.2E-10 | 4.94 (1611,100,88,27) |
| GO:0051276 | chromosome organization | 2.78E-13 | 2.26E-10 | 5.26 (1611,87,88,25) |
| GO:1901990 | regulation of mitotic cell cycle phase transition | 2.99E-13 | 2.21E-10 | 5.01 (1611,95,88,26) |
| GO:0051983 | regulation of chromosome segregation | 2.35E-12 | 1.59E-9 | 7.78 (1611,40,88,17) |
| GO:0045930 | negative regulation of mitotic cell cycle | 9.58E-12 | 6.00E-09 | 5.91 (1611,62,88,20) |
| GO:0045786 | negative regulation of cell cycle | 1.18E-11 | 6.84E-9 | 4.53 (1611,101,88,25) |
| GO:0010948 | negative regulation of cell cycle process | 1.3E-11 | 7.08E-9 | 5.49 (1611,70,88,21) |
| GO:1901988 | negative regulation of cell cycle phase transition | 3.17E-11 | 1.61E-8 | 6.34 (1611,52,88,18) |
| GO:0010965 | regulation of mitotic sister chromatid separation | 4.85E-11 | 2.32E-8 | 9.52 (1611,25,88,13) |
| GO:1905818 | regulation of chromosome separation | 4.85E-11 | 2.19E-8 | 9.52 (1611,25,88,13) |
| GO:0045787 | positive regulation of cell cycle | 5.77E-11 | 2.47E-8 | 5.13 (1611,75,88,21) |
| GO:1901991 | negative regulation of mitotic cell cycle phase transition | 1.69E-10 | 6.87E-8 | 6.22 (1611,50,88,17) |

**Table S5.**
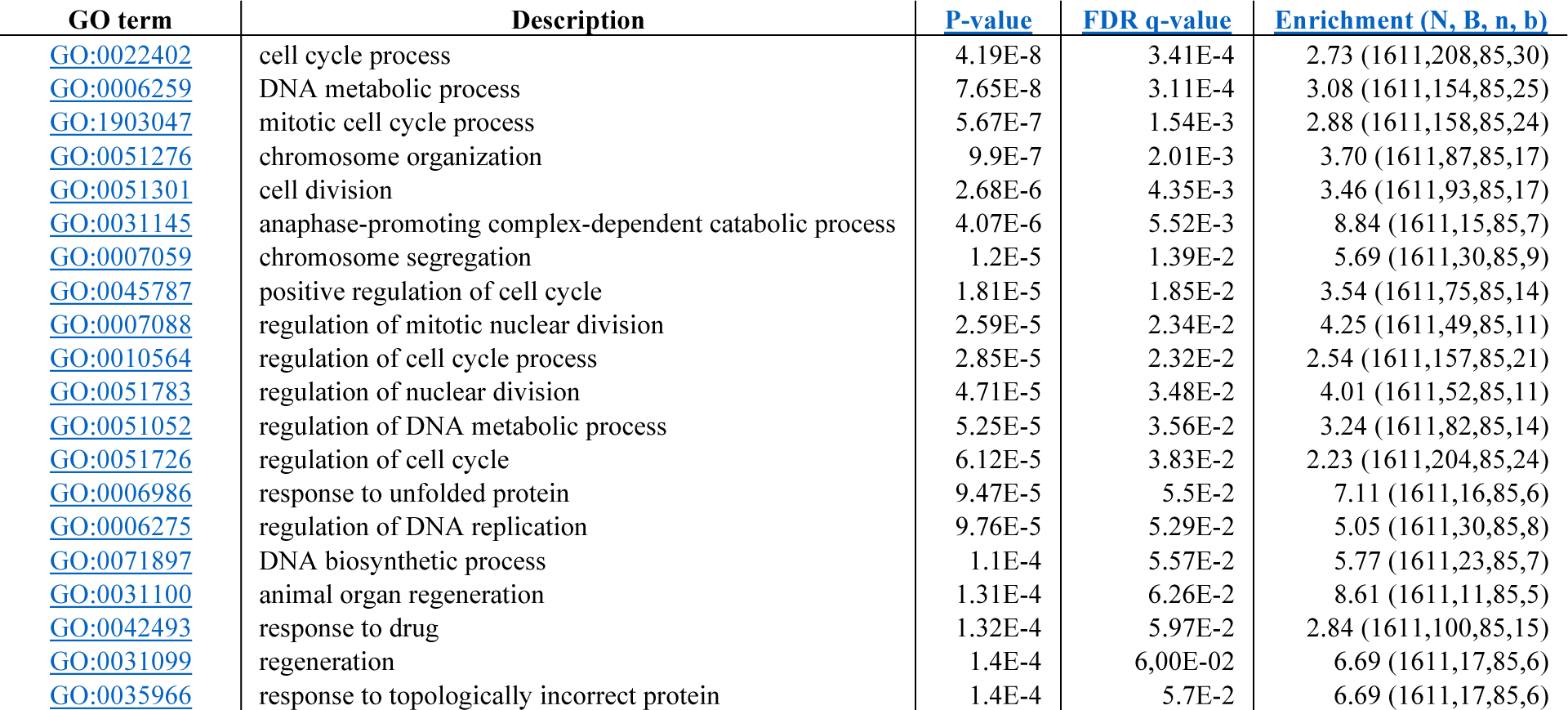
First 20 entries obtained by GO term analysis (*16, 17*) of the weights associated with PC3 are still dominated by the cell cycle though with lower p-values.

**Table S6.** First 20 entries obtained by GO term analysis (*16, 17*) of the weights associated with DC1 are dominated by the cell cycle.

| GO term | Description | P-value | FDR q-value | Enrichment (N, B, n, b) |
| --- | --- | --- | --- | --- |
| GO:0022402 | cell cycle process | 2.03E-27 | 1.65E-23 | 4.48 (1611,208,95,55) |
| GO:1903047 | mitotic cell cycle process | 3.69E-25 | 1.5E-21 | 5.04 (1611,158,95,47) |
| GO:0051726 | regulation of cell cycle | 4.25E-17 | 1.15E-13 | 3.66 (1611,204,95,44) |
| GO:0051301 | cell division | 1.7E-15 | 3.46E-12 | 5.29 (1611,93,95,29) |
| GO:0006259 | DNA metabolic process | 5.2E-14 | 8.46E-11 | 3.85 (1611,154,95,35) |
| GO:0051276 | chromosome organization | 2.2E-13 | 2.99E-10 | 5.07 (1611,87,95,26) |
| GO:0010564 | regulation of cell cycle process | 6.47E-13 | 7.52E-10 | 3.67 (1611,157,95,34) |
| GO:0007346 | regulation of mitotic cell cycle | 6.2E-12 | 6.3E-9 | 3.85 (1611,132,95,30) |
| GO:0044770 | cell cycle phase transition | 4.43E-11 | 4.01E-8 | 5.47 (1611,62,95,20) |
| GO:1901987 | regulation of cell cycle phase transition | 6.06E-11 | 4.93E-8 | 4.24 (1611,100,95,25) |
| GO:0045786 | negative regulation of cell cycle | 7.7E-11 | 5.7E-8 | 4.20 (1611,101,95,25) |
| GO:1902749 | regulation of cell cycle G2/M phase transition | 8.6E-11 | 5.83E-8 | 5.99 (1611,51,95,18) |
| GO:1901990 | regulation of mitotic cell cycle phase transition | 1.26E-10 | 7.9E-8 | 4.28 (1611,95,95,24) |
| GO:0010389 | regulation of G2/M transition of mitotic cell cycle | 1.97E-10 | 1.14E-7 | 6.13 (1611,47,95,17) |
| GO:0044772 | mitotic cell cycle phase transition | 2.12E-10 | 1.15E-7 | 5.37 (1611,60,95,19) |
| GO:0007017 | microtubule-based process | 3.32E-10 | 1.69E-7 | 3.80 (1611,116,95,26) |
| GO:1903046 | meiotic cell cycle process | 1.44E-9 | 6.88E-7 | 7.60 (1611,29,95,13) |
| GO:0010948 | negative regulation of cell cycle process | 4.1E-9 | 1.85E-6 | 4.60 (1611,70,95,19) |
| GO:0006260 | DNA replication | 5.81E-9 | 2.49E-6 | 5.43 (1611,50,95,16) |
| GO:0051983 | regulation of chromosome segregation | 1.55E-8 | 6.33E-6 | 5.94 (1611,40,95,14) |

**Table S7.**
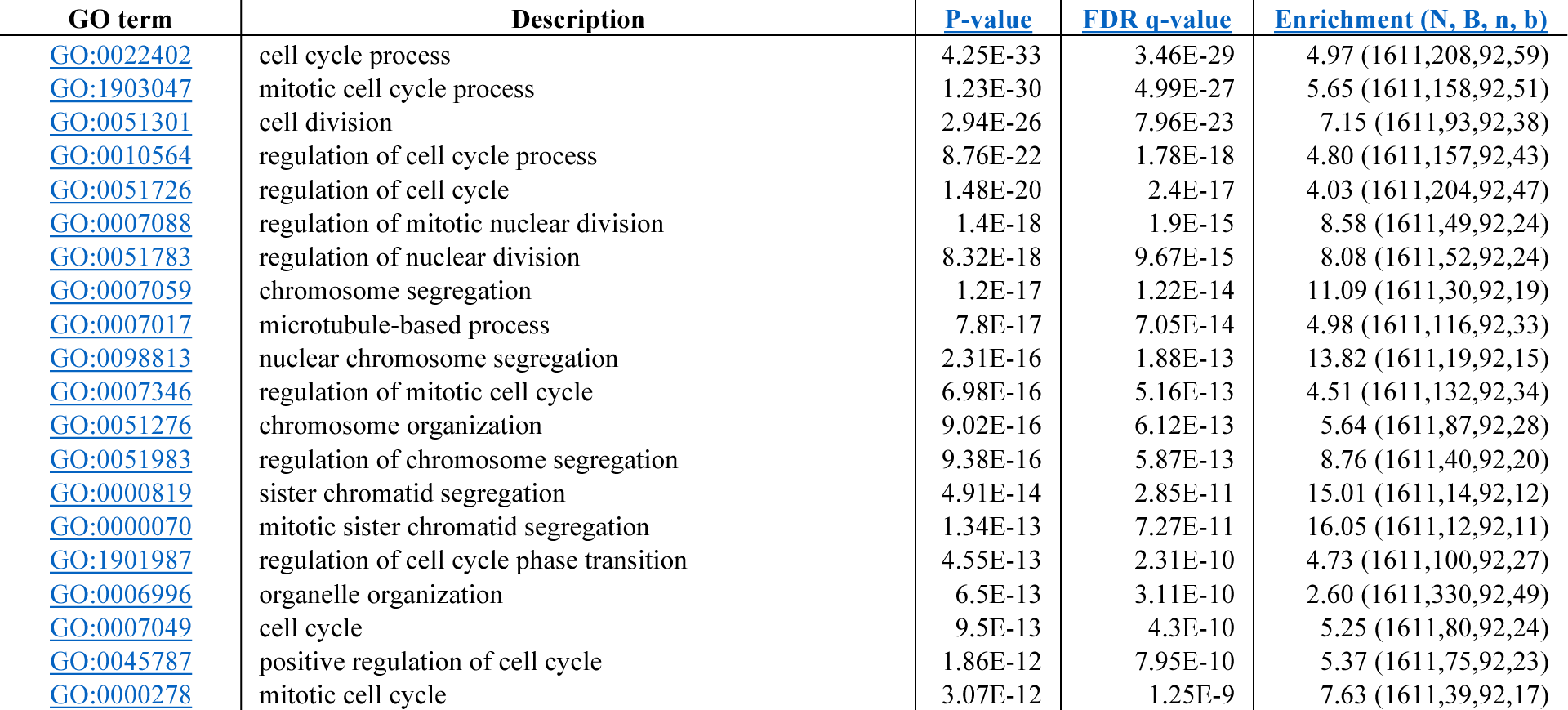
First 20 entries obtained by GO term analysis (*16, 17*) of the weights associated with DC2 are dominated by the cell cycle.

| GO term | Description | P-value | FDR q-value | Enrichment (N, B, n, b) |
| --- | --- | --- | --- | --- |
| GO:0022402 | cell cycle process | 4.25E-33 | 3.46E-29 | 4.97 (1611,208,92,59) |
| GO:1903047 | mitotic cell cycle process | 1.23E-30 | 4.99E-27 | 5.65 (1611,158,92,51) |
| GO:0051301 | cell division | 2.94E-26 | 7.96E-23 | 7.15 (1611,93,92,38) |
| GO:0010564 | regulation of cell cycle process | 8.76E-22 | 1.78E-18 | 4.80 (1611,157,92,43) |
| GO:0051726 | regulation of cell cycle | 1.48E-20 | 2.4E-17 | 4.03 (1611,204,92,47) |
| GO:0007088 | regulation of mitotic nuclear division | 1.4E-18 | 1.9E-15 | 8.58 (1611,49,92,24) |
| GO:0051783 | regulation of nuclear division | 8.32E-18 | 9.67E-15 | 8.08 (1611,52,92,24) |
| GO:0007059 | chromosome segregation | 1.2E-17 | 1.22E-14 | 11.09 (1611,30,92,19) |
| GO:0007017 | microtubule-based process | 7.8E-17 | 7.05E-14 | 4.98 (1611,116,92,33) |
| GO:0098813 | nuclear chromosome segregation | 2.31E-16 | 1.88E-13 | 13.82 (1611,19,92,15) |
| GO:0007346 | regulation of mitotic cell cycle | 6.98E-16 | 5.16E-13 | 4.51 (1611,132,92,34) |
| GO:0051276 | chromosome organization | 9.02E-16 | 6.12E-13 | 5.64 (1611,87,92,28) |
| GO:0051983 | regulation of chromosome segregation | 9.38E-16 | 5.87E-13 | 8.76 (1611,40,92,20) |
| GO:0000819 | sister chromatid segregation | 4.91E-14 | 2.85E-11 | 15.01 (1611,14,92,12) |
| GO:0000070 | mitotic sister chromatid segregation | 1.34E-13 | 7.27E-11 | 16.05 (1611,12,92,11) |
| GO:1901987 | regulation of cell cycle phase transition | 4.55E-13 | 2.31E-10 | 4.73 (1611,100,92,27) |
| GO:0006996 | organelle organization | 6.5E-13 | 3.11E-10 | 2.60 (1611,330,92,49) |
| GO:0007049 | cell cycle | 9.5E-13 | 4.3E-10 | 5.25 (1611,80,92,24) |
| GO:0045787 | positive regulation of cell cycle | 1.86E-12 | 7.95E-10 | 5.37 (1611,75,92,23) |
| GO:0000278 | mitotic cell cycle | 3.07E-12 | 1.25E-9 | 7.63 (1611,39,92,17) |

**Fig. S1.**
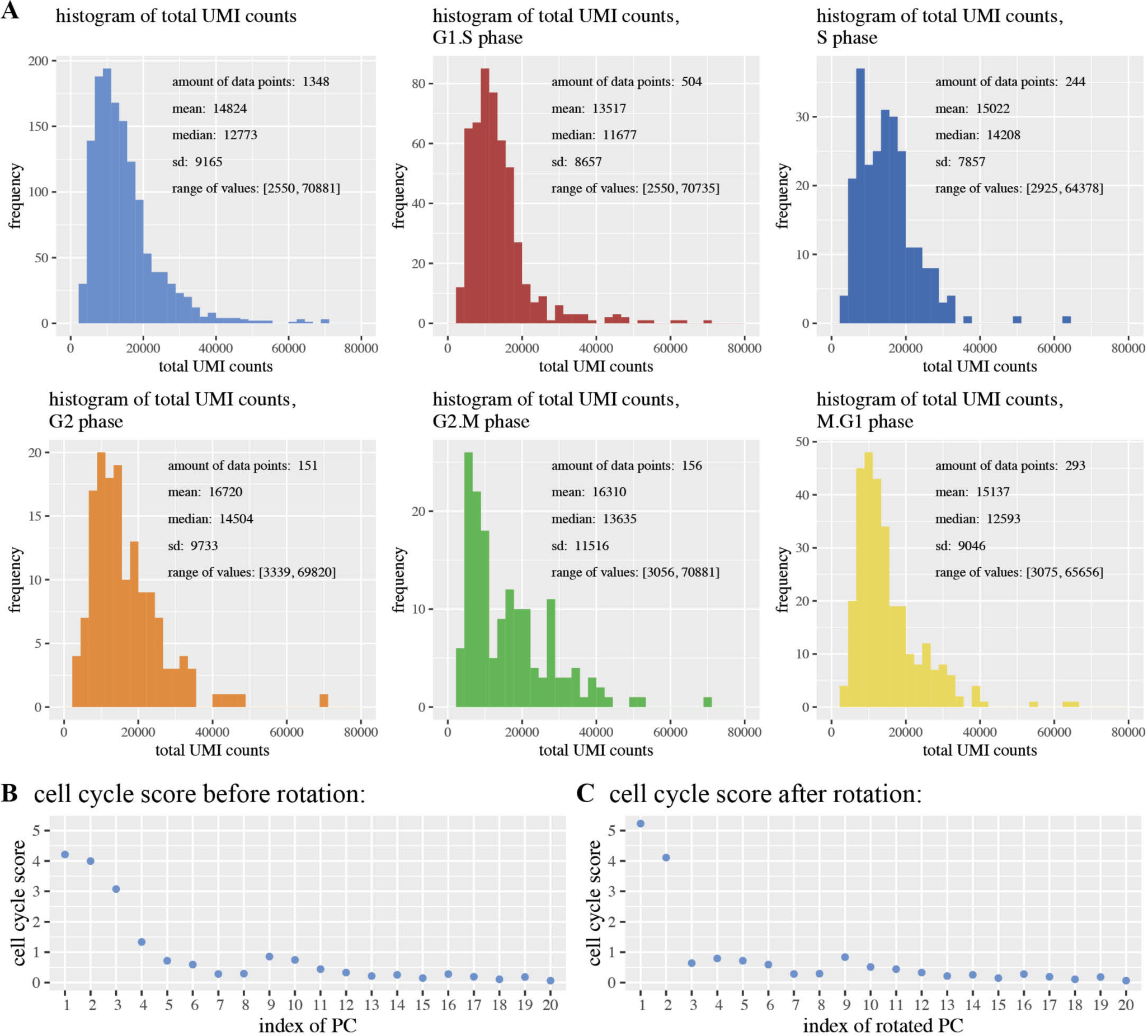
Additional statistics of HeLa data set 1. (**A**) Distribution of total UMI counts per cell for each cell cylce phase for data set 1. (top left) All data points to be examined, (top middle) only cells placed in G1.S phase, (top right) S phase, (bottom left) G2 phase, (bottom middle) G2.M phase, (bottom right) M.G1 phase. (**B**) Cell cycle scores for the first 20 PCs. (**C**) Cell cycle score for the first 20 rotated PCs. PCs involved in the rotation: PC1, PC2, PC3, PC4, PC9, PC10. The rotation of PC space entails the first two components only to have significant cell cycle scores, suggesting that additional components are free of cell cycle effects.

**Fig. S2.**
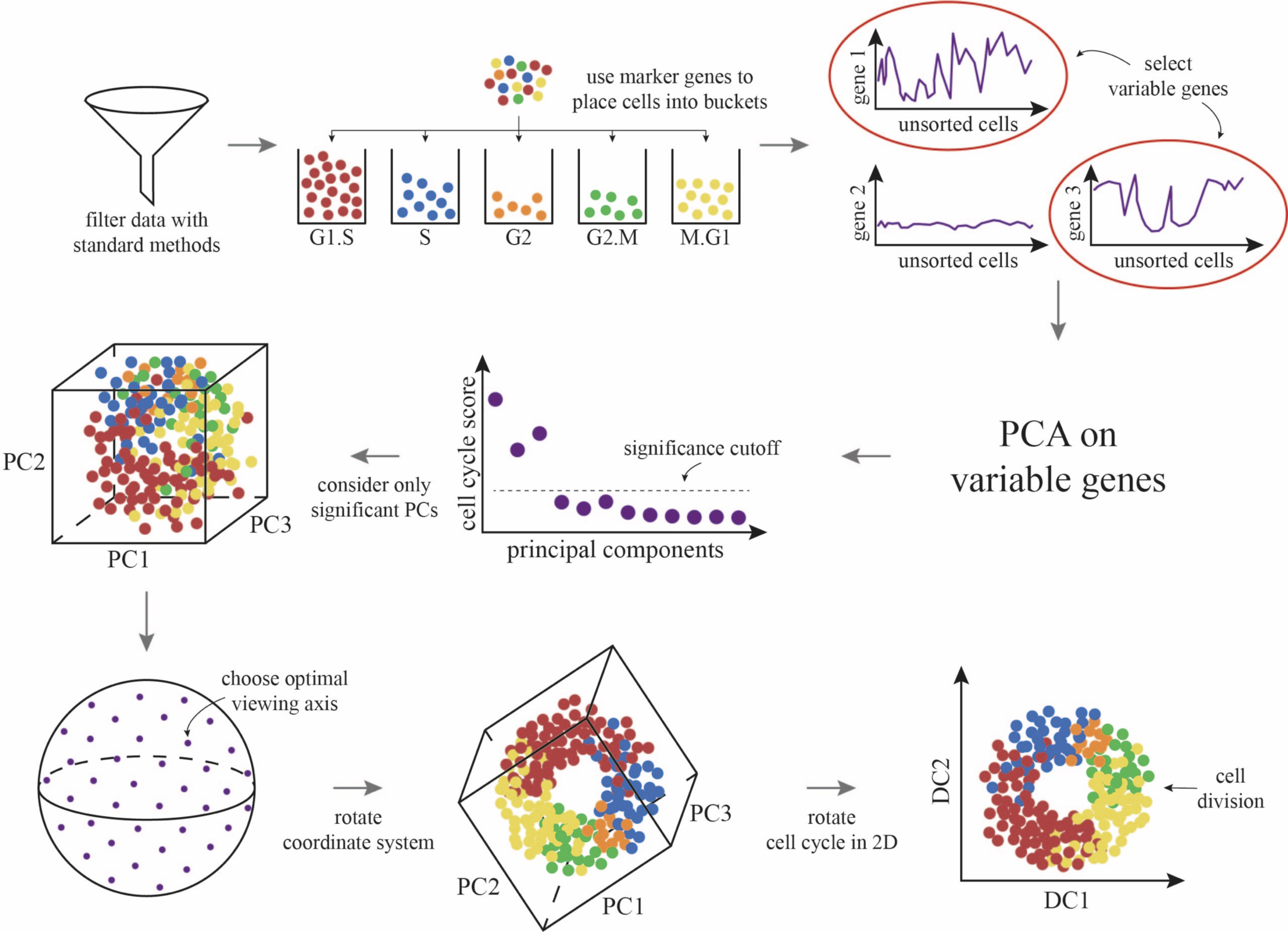
The principle steps of our algorithm REVELIO for extracting the cell cycle from the data. After the data is filtered by standard methods, we divide the cells into buckets with the help of marker genes (*8, 10*). Next, we select variable genes (*11*) and apply PCA on the reduced data set. Afterwards, we utilize a cell cycle score (Methods) to judge which PCs are influenced by the cell cycle. The significant PCs are used to construct three-dimensional subspaces. We then choose an optimal viewing axis by minimizing the cell cycle score along the viewing axis (Methods). The coordinate system is rotated linearly and the cell cycle is obtained only within the plane spanned by the first two new axes (DC1, DC2). Within the DC1-DC2-plane, we estimate the time point of cell division and rotate the cell cycle plane accordingly (Methods).

**Fig. S3.**
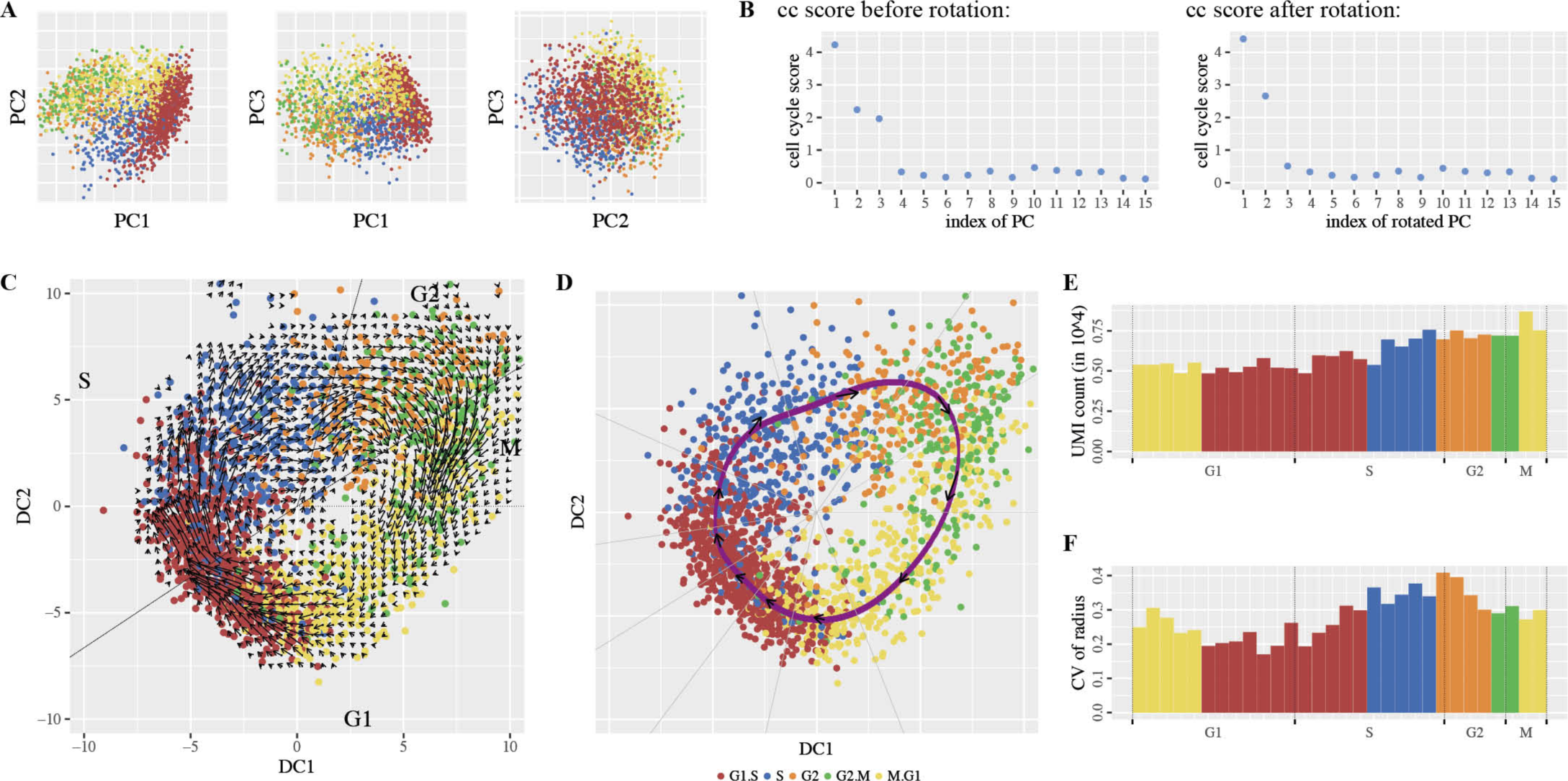
Data set 2 (HeLa cells) confirms that a two-dimensional annulus represents the cell cycle in HeLa cell populations. This data set has almost twice as many cells as the previous data set 1 at less than half the sequencing depth. (**A**) Pairwise PC plots again indicate the dynamical components to be slanted with respect to the original PCs. A clear periodic structure is completely absent. (**B**) (left) Again the cell cycle score concurs with our observations from the PC plots. (right) The algorithm manages to place all cell cycle effects w.r.t. clustering within the first two components. (**C**) After rotation and the overlaying of RNA velocity, a clear cell cycle in the form of an annulus is observed. Motion of the cells appears strongest during G1-S and M phase similar to the data set 1. (**D**) The average RNA velocity per interval is clearly tangential to the trajectory of an averaged cell. (**E**) The increase of average total UMI count per interval and the sharp drop between last and first bins are consistent with our previous observations. (**F**) Similarly, the dynamics of the CV of the radius along the cell cycle resemble our observations from fig. S7, exhibiting a peak around the S-G2 transition and a drop towards M phase.

**Fig. S4.**
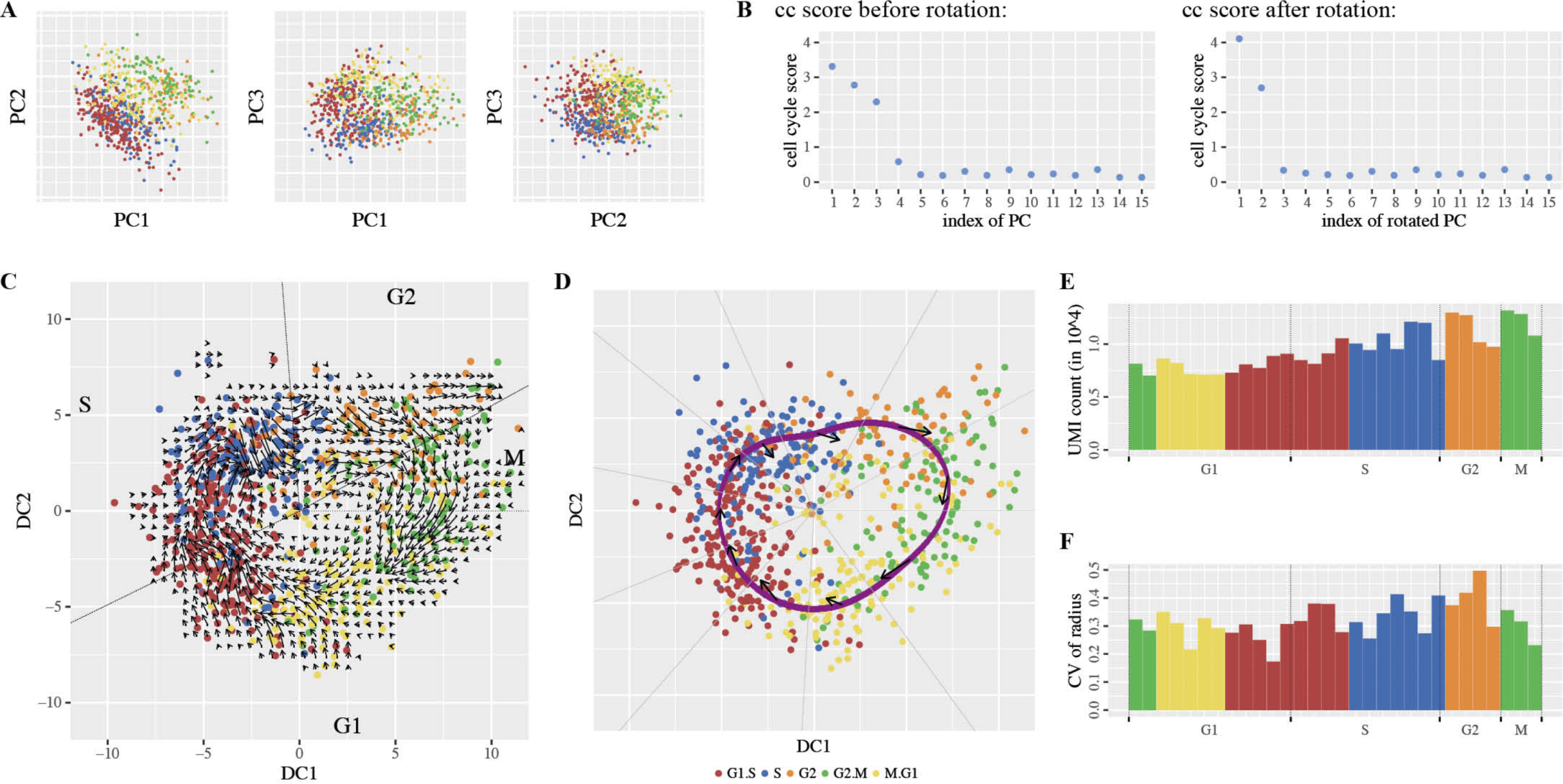
Data set 3 (HEK cells) confirms that our observations and the algorithm are transferable to a different cell type. (**A**) Pairwise PC plots show clustering within the first three components but are very noisy and lack periodic structures. (**B**) The cell cycle score confirms our observations and is in line with previous data sets. (**C**) Rotation of the space once again yields a clear periodic object with clustering according to inferred cell cycle phases. RNA velocity confirms the motion of the cells. The clearest motion again appears around the time point G1-S and throughout M phase. (**D**) While the average velocity on the average cell is mostly tangential, we notice that during S phase and before G1-S, the direction of movement is more directed towards the center of the cycle. (**E**) The drop by factor 1/2 is less clear than what we saw in HeLa data but an increase in UMI counts towards M phase is nevertheless present. We suspect the data to be noisier since we operate with half as many cells as data set 1 and with 2/3 of the sequencing depth. (F) An increase in CV now takes place during G2 phase.

**Fig. S5.**
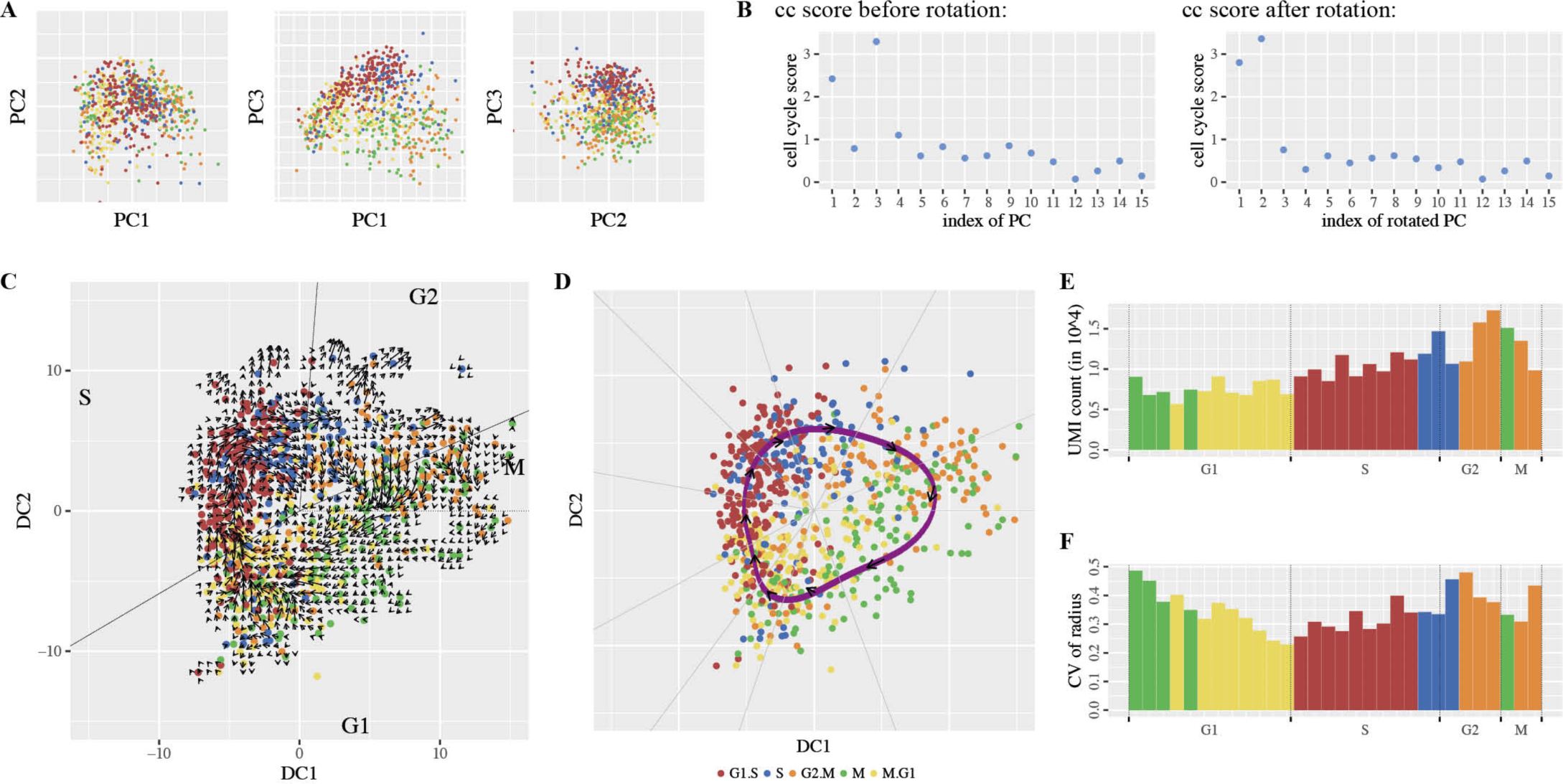
The cell cycle properties appear to be preserved across species as the cell cycle in mouse 3T3 cells is also shown to form a cyclic object in two dimensions. (**A**) Similar to the human cell lines we have investigated previously, pairwise PC plots show clustering w.r.t. cell cycle phases but are not periodic. (**B**) The cell cycle score of PCs and rotated PCs show similar patterns as before. (**C**) Although, we find a lot of noise at our estimated G1-S time point, the rest of the cell cycle in two dimensions is remarkably clear in clustering according to computationally inferred cell cycle phases. The distinctions about variability in velocity magnitude are less clear than before due to high amounts of suspected noise. (**D**) The average RNA velocity on an average individual cell points surprisingly well into the tangential direction. (**E**) Similar to the HEK data set, the drop in average total UMI counts along the cell cycle is less clear and appears to happen slightly earlier than previous data sets. As before, we suspect additional noise incorporated into the data set due to the shallower sequencing depth to be the cause for this. (**F**) The CV of the radius has a peak around the blue and orange clusters within G2 phase. This suggests that the synchronization to cell division might be slightly inaccurate. We also observe a severe peak of the CV during M-G1 transition. Considering the two-dimensional plot in (c) and (d) the large number of outliers around this time point seem to cause this which might be due to noise.

**Fig. S6.**
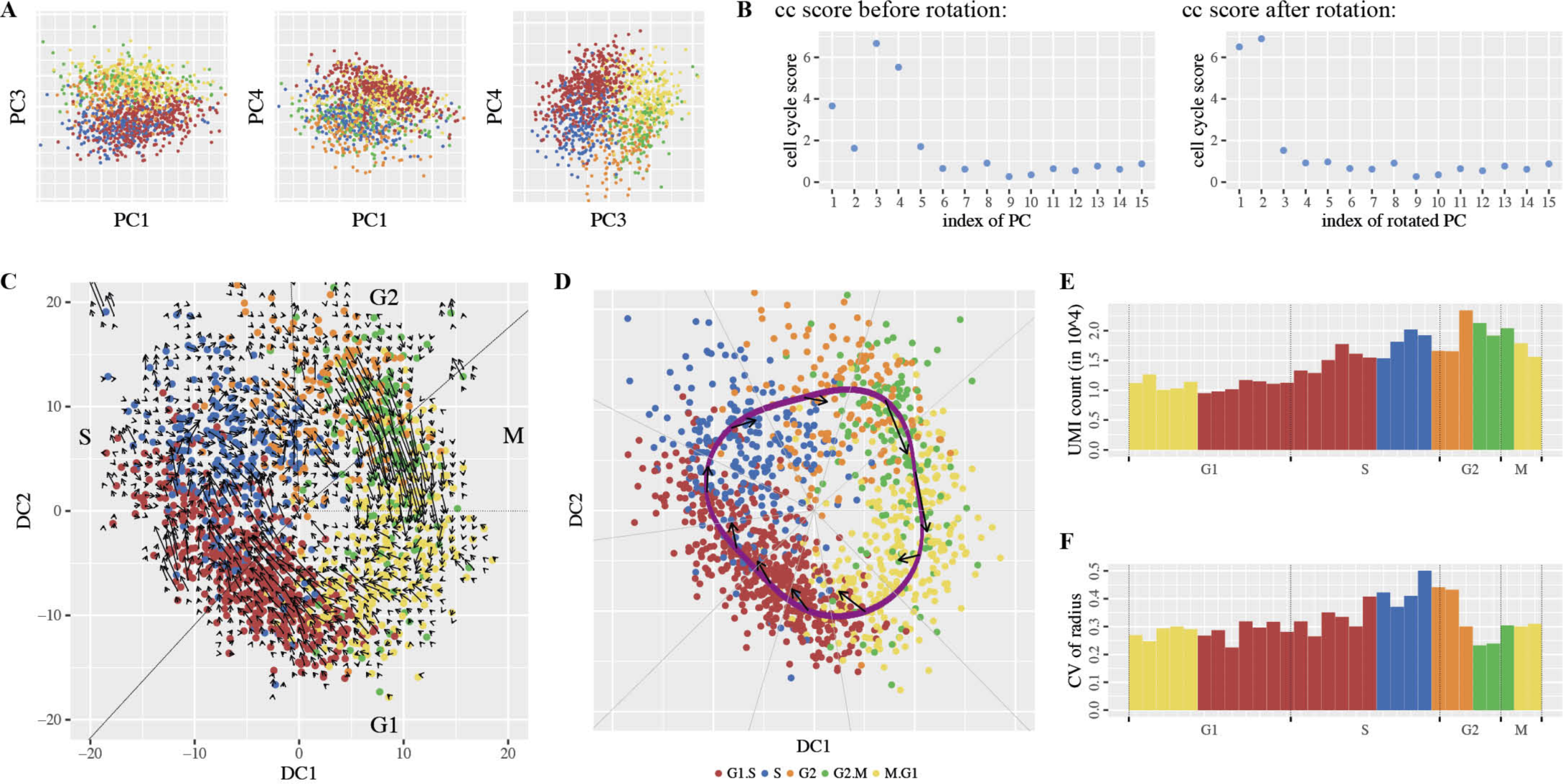
HeLa data set 1.2 utilizing more than 10.000 genes yields two-dimensional cycle after rotation of PCA. The same data as in Fig. 1–3 is analyzed again utilizing all genes for which exonic and intronic reads were measured. (**A**) Pairwise PC plots of PC1, PC3, PC4. PC3 and PC4 on their own already form an acceptable cell cycle. (**B**) (left) The cell cycle score already points to PC3 and PC4 containing most cell cycle effects. (right) After rotation, all cell cycle effects are contained in DC1 and DC2. Additional dimensions exhibit no significant cell cycle score. (**C**) By rotation and overlaying RNA velocity, the cell cycle becomes much clearer and forms an annulus in two dimensions. Our analysis from Fig. 3A holds as the strongest average motion of cells takes place during G1-S and M phase. (**D**) An average spline is added (magenta) which we interpret as an average cell trajectory. The average RNA velocity is approximately tangent to the cyclic trajectory. (**E**) Progression of average total UMI count per interval along the cell cycle shows a sharp drop between the last and first bin, where cell division is suspected to take place. (**F**) CV of the radius increases at the end of S phase and drops sharply during G2 phase before M phase.

**Fig. S7.**
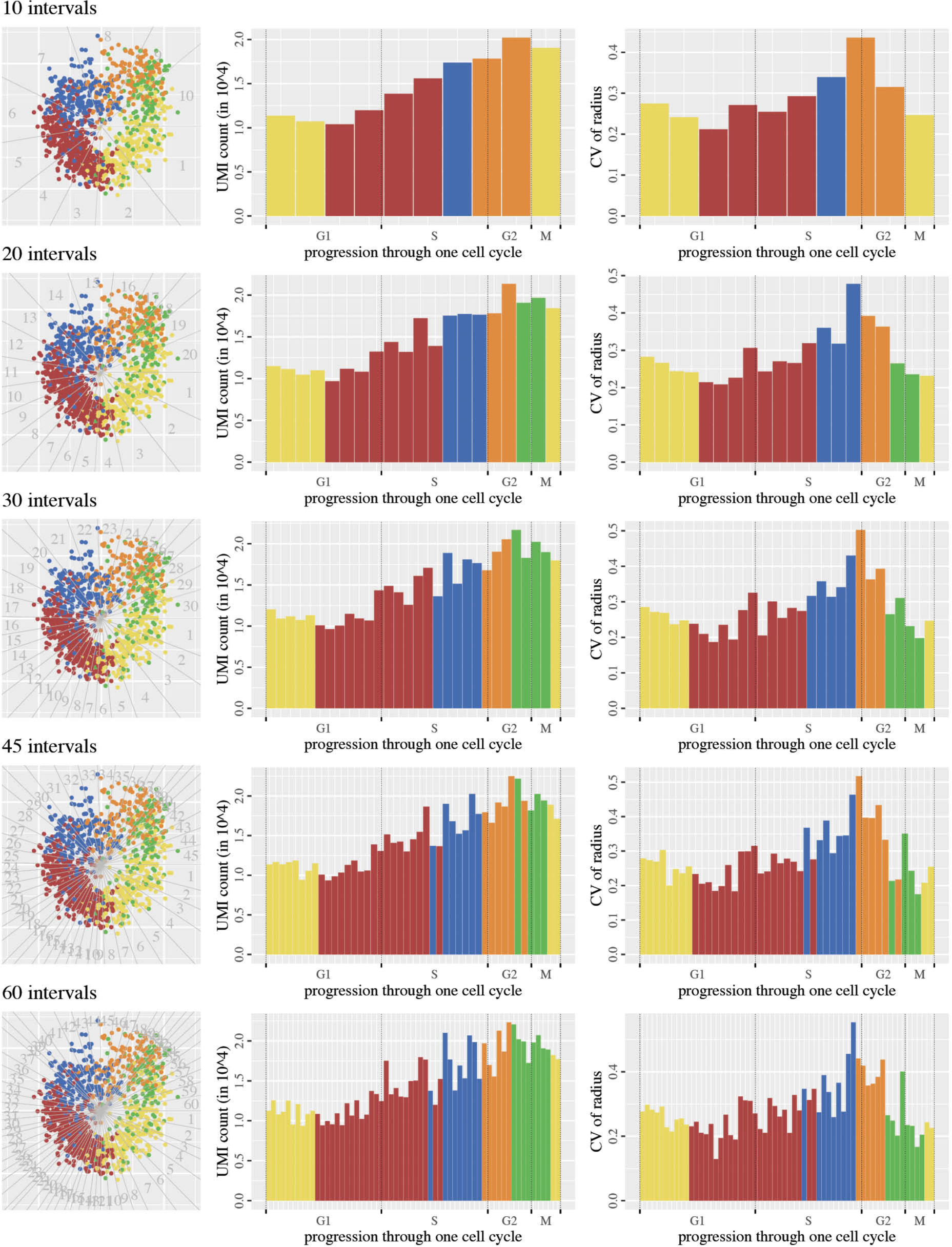
The drop of total UMI count by a factor 1/2 is present within the data independently of the number of bins chosen while the CV of the radius of cells indicates higher variability of cells during the transition S-G2. Each row corresponds to a different binning with increasing number of intervals. Intervals are always chosen to contain the same amount of cells. The left column indicates what the dissection of the cell cycle looks like. The middle column shows the average total amount of UMIs per cell contained within each interval as was already shown in Fig. 2A, bottom. We observe that the drop by factor 1/2 we saw previously is present within any binning. The right most column portrays the coefficient of variation of the radius of cells taken from the two-dimensional cell cycle plot. We observe a consistent peak towards the end of S and beginning of G2 phase and a slight increase of the CV during the beginning of G1 phase. This lines up well with increased variability of the direction of velocity in Fig. 3A, providing another indication that cells during those phases are more variable in their gene expression and are more tightly regulated when entering M phase where the CV drops sharply.

**Fig. S8.**
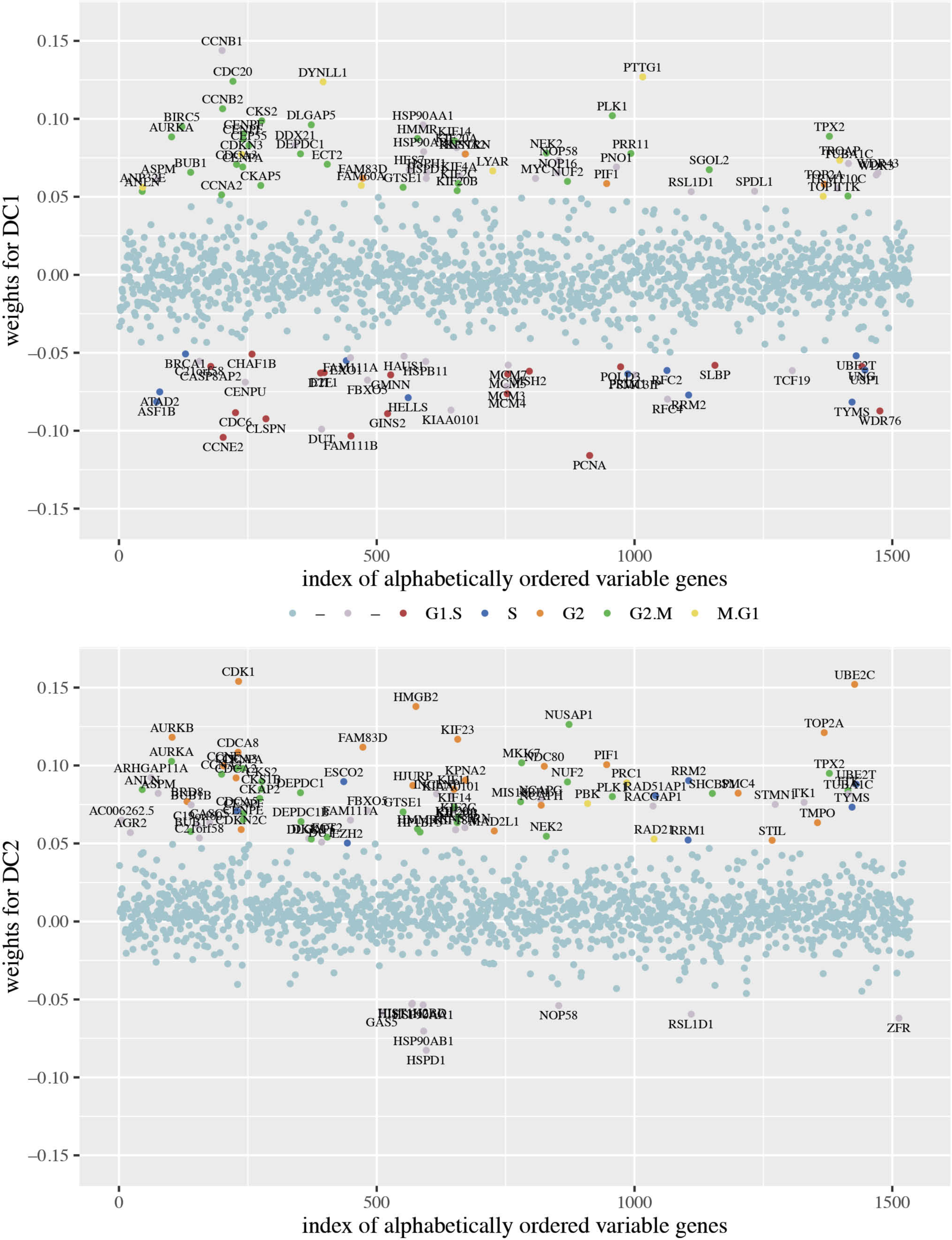
Genes associated with dynamical components mostly correspond to known oscillating genes. (top) Weights of genes that span DC1. Colors indicate if a gene is known to be oscillating from Whitfield et al. (*10*). Negative values (corresponding to the left part of the x-axis of Fig. 1C) are mostly associated with G1-S and S while positive values (right part of x-axis in Fig. 1C) correspond to M phase. (bottom) Weights of genes that span DC2. Positive values are associated with the transition S-G2 and M phase. Very few genes have significant negative weights for DC2. Within our cell cycle from Fig. 1C the lower part of the y-axis corresponds to G1 phase. Thus, this plot confirms that almost no variable genes are active during G1 phase making it difficult to classify cycling cells into G1 because of the lack of marker genes.

**Fig. S9.**
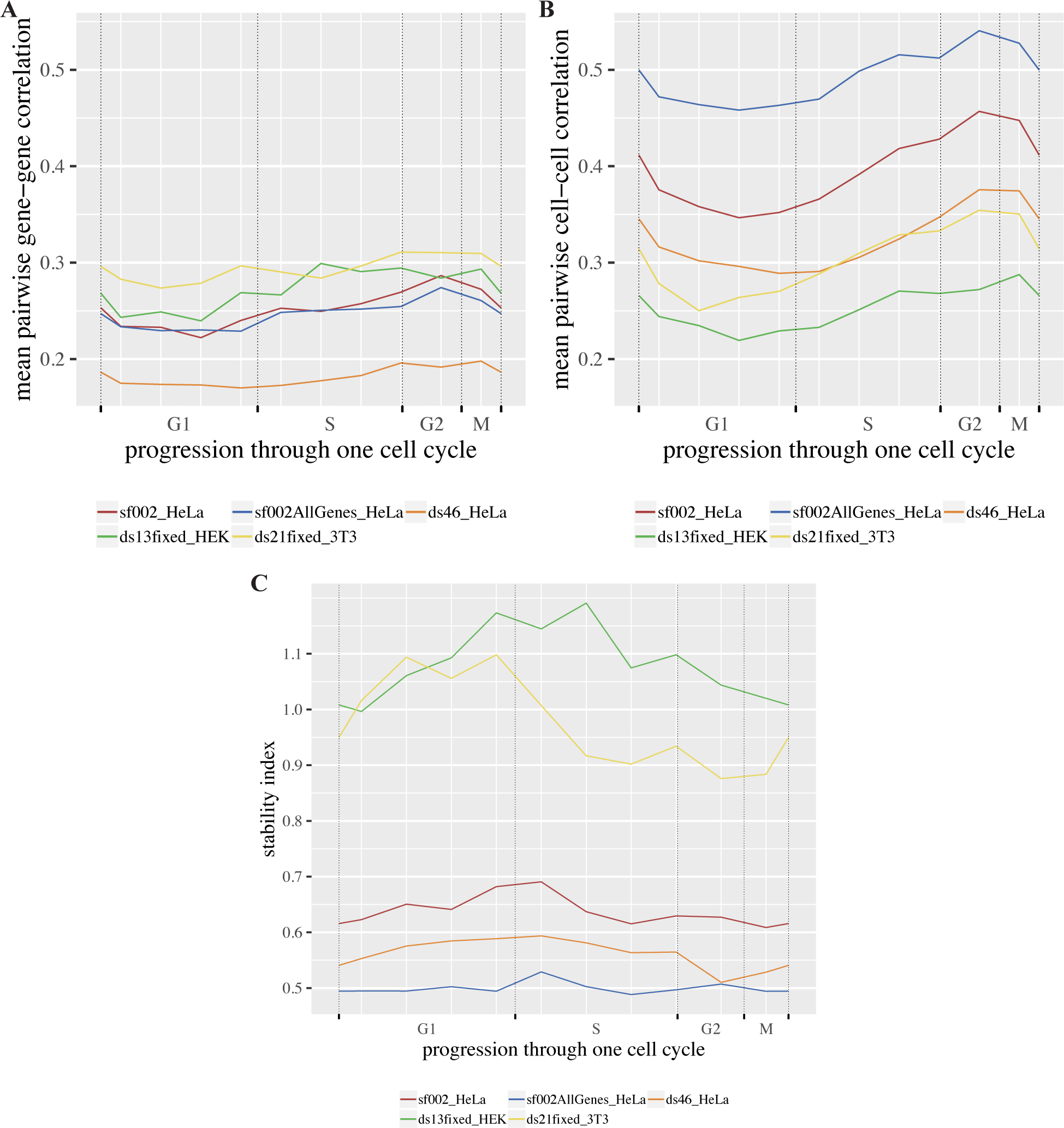
Stability index (*22*) along the cell cycle does not indicate a critical state transition. (**A**) The average pairwise gene-gene correlation increases slightly from G1 to G2 phase. (**B**) We observe a clear trend for average pairwise cell-cell correlations along the cell cycle with an increase after cells transition to S phase, reaching its peak at the end of G2 phase, followed by a relatively sharp drop during M phase and through cell division. It appears that in preparation of M phase, cells are getting more and more coordinated in their gene expression while during G1 phase, gene expression is least correlated. (**C**) The resulting stability index (ratio of gene-gene to cell-cell correlations) (*22*) indicates a rather homogeneous stability, in particular within our HeLa data sets. The HEK and 3T3 data are the noisier data sets and drawing consistent meaning from their graphs is more difficult. The stability index exhibits very similar values throughout the cycle, because we observe simultaneous increase of cell-cell and gene-gene correlations along the cell cycle. As a critical state transition is accompanied by a rise of the stability index, we suspect that such a transition is not taking place along the cell cycle of immortalized cell lines.

**Fig. S10.**
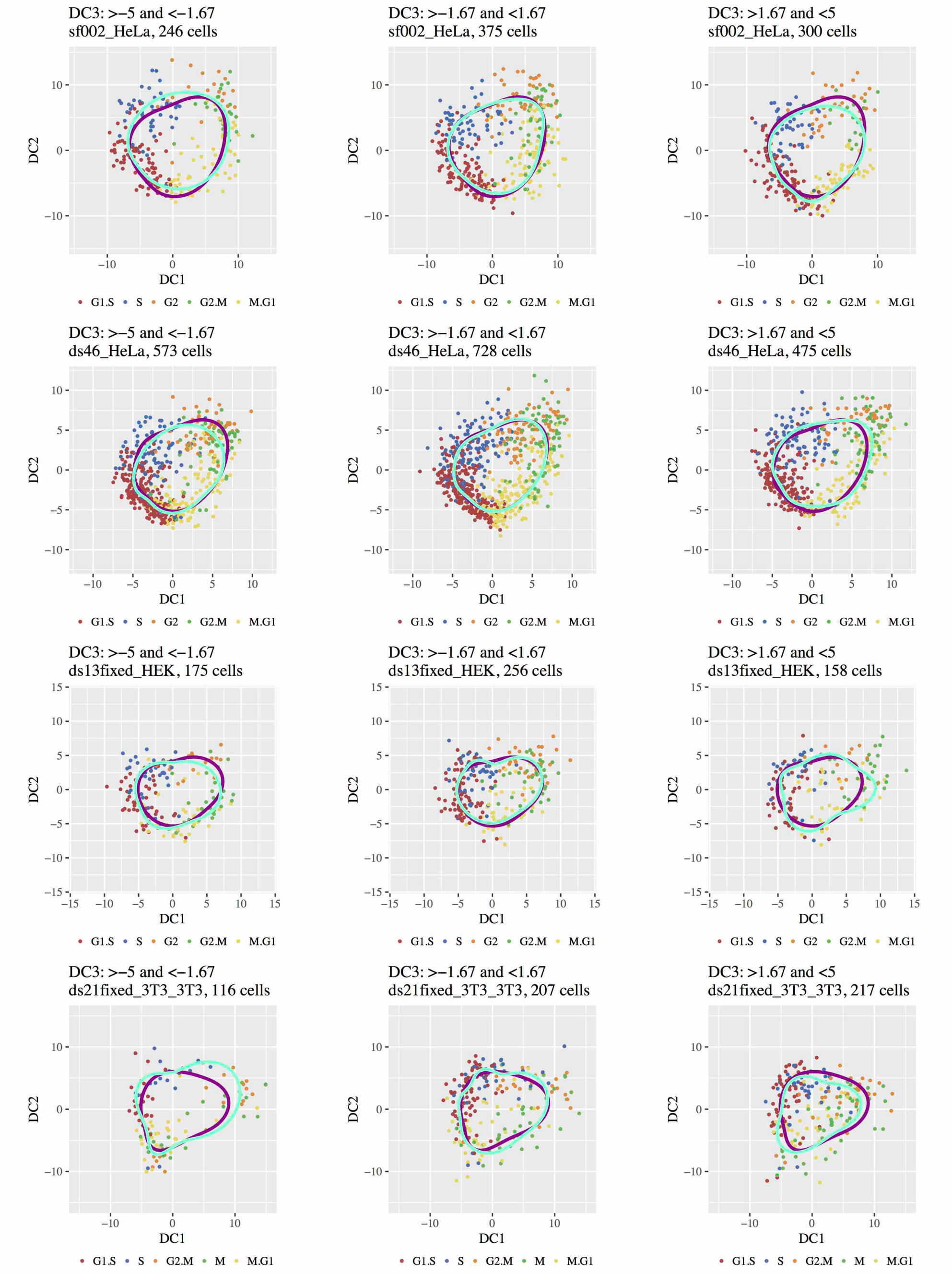
Average trajectories for different height ranges of the cylinder in four data sets. The cylinder formed by the data cloud in DC1-DC2-DC3 is divided into three subsections along the cylinder axis (DC3). Each row corresponds to a different data set and each column signifies a different cylinder slice. In purple the average cell cycle of the respective population is displayed, whereas in light blue the average cell cycle trajectory of all cells displayed in a particular plot is shown. We observe that the average trajectories do not significantly differ from the population average. This implies that a cell’s placement along the cylinder axis does not influence gene expression of the cell cycle.

